# Dual domain recognition determines SARS-CoV-2 PLpro selectivity for human ISG15 and K48-linked di-ubiquitin

**DOI:** 10.1101/2021.09.15.460543

**Authors:** Pawel M. Wydorski, Jerzy Osipiuk, Benjamin T. Lanham, Christine Tesar, Michael Endres, Elizabeth Engle, Robert Jedrzejczak, Vishruth Mullapudi, Karolina Michalska, Krzysztof Fidelis, David Fushman, Andrzej Joachimiak, Lukasz A. Joachimiak

## Abstract

The Papain-like protease (PLpro) is a domain of a multi-functional, non-structural protein 3 of coronaviruses. PLpro cleaves viral polyproteins and posttranslational conjugates with poly-ubiquitin and protective ISG15, composed of two ubiquitin-like (UBL) domains. Across coronaviruses, PLpro showed divergent selectivity for recognition and cleavage of posttranslational conjugates despite sequence conservation. We show that SARS-CoV-2 PLpro binds human ISG15 and K48-linked di-ubiquitin (K48-Ub_2_) with nanomolar affinity and detect alternate weaker-binding modes. Crystal structures of untethered PLpro complexes with ISG15 and K48-Ub_2_ combined with solution NMR and cross-linking mass spectrometry revealed how the two domains of ISG15 or K48-Ub_2_ are differently utilized in interactions with PLpro. Analysis of protein interface energetics predicted differential binding stabilities of the two UBL/Ub domains that were validated experimentally. We emphasize how substrate recognition can be tuned to cleave specifically ISG15 or K48-Ub_2_ modifications while retaining capacity to cleave mono-Ub conjugates. These results highlight alternative druggable surfaces that would inhibit PLpro function.

## INTRODUCTION

The COVID-19 pandemic is caused by Severe Acute Respiratory Syndrome Coronavirus 2 (SARS-CoV-2) from the *Coronaviridae* family^1,2^, a spherical, enveloped, non-segmented, (+) sense RNA virion with a ∼30 kbs genome. The RNA is used for synthesis of two polyproteins (Pp1a/Pp1ab), which are processed by two viral proteases: papain-like protease (PLpro) and 3C-like protease (Mpro) and assembly of the replication-transcription complex (RTC). PLpro is a domain of non-structural protein 3 (Nsp3). It cleaves three sites in SARS-CoV-2 polyproteins yielding Nsp1, Nsp2 and Nsp3. The “LXGG↓XX” motif found in the polyproteins is essential for protease recognition and cleavage. PLpro has been shown to have additional functions, including the best characterized deubiquitinating^4–6^ and deISGylating activities^7,8^, occurring by the cleavage of conjugates having the “LRGG” sequence motif found at the C-terminus of ubiquitin (Ub) and ISG15. These activities dysregulate Ub- and ISG15-dependent pathways which play important roles in protein degradation, vesicular trafficking, inflammatory, anti-pathogen responses and homeostasis^8,9^. Interestingly, the SARS-CoV-1 PLpro has a strong preference for hydrolysis of K48-linked polyUb conjugates, in contrast to the MERS-CoV enzyme that acts as a “general” broad-specificity deubiquitinase (DUB)^10^. Removing these modifications disturbs interferon (IFN) expression and blocks NF-kappaB signaling^10^, and cleaving off ISG15 from STAT induces up-regulation of TGF-β1^11^. Deubiquitination and deISGylation were shown to be utilized by a broad family of viruses that include Coronaviruses, Hepadnaviruses, Nairoviruses, Arterioviruses, Picornaviruses and Aphthoviruses^12^. Some other PLpro functions involve direct cleavage of host proteins influencing wide-ranging processes from blood coagulation to nuclear transport^13–15^. PLpro may also play roles beyond its proteolytic activity^6^, illustrating its diverse and complex functions^3^. The SARS-CoV-2 PLpro (PLpro^CoV-2^) sequence and structural fold are conserved among SARS-CoV-1 (83% identical), MERS-CoV (30% identical) and other coronaviruses. Despite low sequence identities (∼10%)^7^, PLpros also share common structural architecture and catalytic site with the human ubiquitin specific proteases (USPs), one of the five distinct deubiquitinating enzyme (DUB) families. Interestingly, one of these USPs (USP18) is specific for cleaving off ISG15 conjugates in humans and other vertebrates, and PLpro^CoV-2^ deISGylation activity may potentiate USP18 regulatory function^16^.

The PLpro^CoV-2^ differs significantly from Mpro, which was shown to recognize linear sequence motif^17^, as it encodes proteolytic activities that control viral polyprotein cleavage but also is processing conjugates of host and viral proteins including polyUb and ISG15. Therefore, PLpro^CoV-2^, in addition to recognizing “LXGG↓XX” linear motif, can also specifically interact with protein surfaces presented in three-dimensional structures and using this mechanism can recognize and select different substrates. Decoupling the basal polyprotein cleavage activity from those required to process polyUb or ISG15 is important to understand their role in viral pathogenesis, how these activities influence interaction of the virus with the host immune system and how changes in these interactions modify host antiviral responses and influence disease outcomes.

Ubiquitination is an essential post-translational modification (PTM) engaged in multiple functions in humans, including signaling and proteasome-dependent protein degradation^18^. The modification is mediated by the Ub-conjugating system and could be reversed by DUBs^8^. ISG15 is an IFNα-stimulated gene that is a critical component of the antiviral response^19,20^. ISG15 is processed and subsequently activated in a manner similar to Ub using interferon-induced factors that follow the ubiquitination-like E1, E2 and E3 enzyme cascade to mediate co-translational ISGylation – an addition of ISG15, via its C-terminal LRGG motif, to substrate lysine residues^21^. It is not precisely clear how ISG15 interferes with viral processes but it is believed that tagging newly translated viral proteins with ISG15 sterically prevents their folding, assembly or interactions^22^. The level of ISGylation is controlled by interferon and USP18^16^. The free unconjugated ISG15 form can exist intracellularly or be secreted to function as a cytokine, linked to the induction of a cytokinin storm^23,24^. Removal of Ub and ISG15 conjugates from specific substrates in host cells may have a diverse impact on numerous cellular processes and specifically may disrupt the host response to viral infection^19,20,25^. PLpro can recognize both appendages and cleave them off as they share PLpro recognition motif at their C-termini. PolyUb and ISG15 have common other structural features: ISG15 comprises two Ub-like (UBL) domains and mimics a head-to-tail linked di-ubiquitin (Ub_2_). While K48-linked Ub_2_ (K48-Ub_2_) and ISG15 are similar both in sequence and fold, the topologies of how the two domains are linked are distinct. How PLpro discriminates between different ubiquitin linkage types and specifically between Ub_2_ and ISG15 substrates is still unknown. Understanding how PLpro discriminates between different substrates will help uncover how these additional proteolytic activities contribute to viral pathogenesis.

Recently published work suggested that mutations in PLpro^CoV-1^ to PLpro^CoV-2^ changed its binding preference from K48-Ub_2_ to human ISG15 (hISG15)^26,27^. Inspired and attracted by these results we investigated interaction of PLpro^CoV-2^ with mono-, di-, and tri-ubiquitins (Ub_1_, Ub_2_, Ub_3_) and hISG15. We employed complementary biochemical, structural X-ray, NMR, mutagenesis and computational approaches, to understand how PLpro^CoV-2^ can differentiate between linear recognition motifs, Ub_1_, K48-Ub_2_ and hISG15 substrates. We find that PLpro^CoV-2^ binds both hISG15 and K48-Ub_2_ with high and similar affinity but shows weaker interactions with Ub_1_. We also find lower affinity alternate binding modes for hISG15, K48-Ub_2_ and Ub_1_ which can be explained by non-stoichiometric binding modes and sequence preference outside the conserved recognition motif. We also observe how amino acid substitutions in the first position <u>**X**</u> following G in the “LXGG↓(<u>**X**</u>)” motif impact peptide cleavage. To reveal details of substrate recognition, we determined structures of non-covalent complexes of PLpro^CoV-2^ (single C111S and double C111S,D286N proteolytically inactive mutants) with hISG15 and K48-Ub_2_. These are the first crystal structures of non-modified, complete complexes and they uncover that hISG15 binding is determined by recognition of both UBL domains while K48-Ub_2_ is recognized mainly through the proximal Ub. We further examined the PLpro binding to K48-Ub_2_, Ub_1_ and hISG15 using NMR experiments. These data, together with cross-linking mass spectrometry (XL-MS) and X-ray crystallography suggest that PLpro^CoV-2^ interacts with both UBL domains of ISG15 whereas K48-Ub_2_ is recognized largely through the proximal Ub, with distal Ub contributing to binding through alternative interactions. We used modelling to predict alternative modes of PLpro binding to substrates that are consistent with cross-linking data. Finally, we tested our binding models by performing an *in silico* ΔΔG alanine scan on PLpro^CoV-2^ in complex with K48-Ub_2_/hISG15 substrates and experimentally validated their binding effects to show differential domain utilization by the PLpro for the two substrates. Our findings uncover binding heterogeneity in PLpro interactions with hISG15 and ubiquitin substrates that decouples binding affinity from the enzyme proteolytic activity.

## RESULTS

### Sequence and topological differences between hISG15 and K48-Ub_2_

Recent biochemical binding and cleavage assays have shown that PLpro^CoV-1^ prefers K48-Ub_2_ while the related PLpro^CoV-2^ binds more tightly to both human and mouse ISG15 (hISG15 and mISG15). A nearly 20-fold higher affinity compared to K48-Ub_2_ suggests that the sequence variation at the substrate binding interface between PLpro^CoV-1^ and PLpro^CoV-2^ may dictate substrate specificity^26,27^. Importantly, these previous studies on PLpro^CoV-1^/PLpro^CoV-2^ binding to Ub_2_ used a non-hydrolysable synthetic triazole linker between the Ubs rather than a native isopeptide K48 linkage, raising questions how linker geometry and rigidity may influence binding to PLpro^26–28^. When considering both domains in K48-Ub_2_ and hISG15, they are 33% identical in sequence (Supplementary Fig. 1a) while the distal (N-terminal) UBL domain of hISG15 is 29% identical to Ub, and the proximal (C-terminal) UBL domain of hISG15 has a slightly higher sequence identity of 37%. hISG15 and mISG15 are 63% sequence identical, and both have similar sequence identities to Ub (Supplementary Fig. 1a, b). Intriguingly, however, a recent study reported that hISG15 binding to PLpro^CoV-2^ has an order of magnitude higher on- and off-rates than mISG15 binding^26^.Importantly, Ub and UBLs of hISG15 and mISG15 also vary in the binding surfaces (Supplementary Fig. 1b, c). The protein domains in hISG15 and K48-Ub_2_ have homologous folds but their sequences and topologies of how the two domains are linked are different (Fig. 1a and Supplementary Fig. 1d).

**Figure 1.**
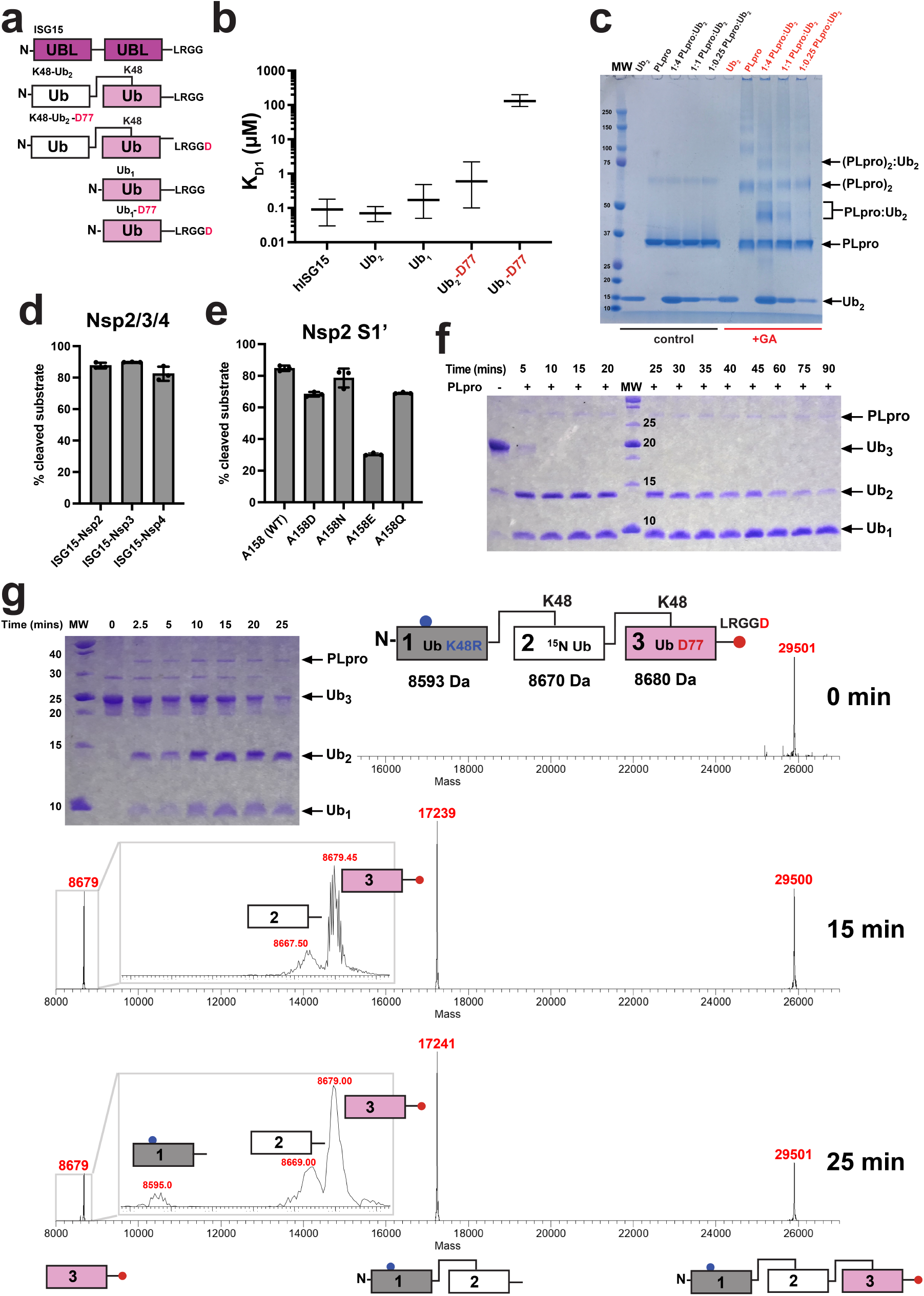
PLpro^CoV-2^ substrate binding and recognition. (**a**) Topology of utilized substrates: ISG15, K48-linked Ub_2_ ending with G76 on the C-terminus (K48-Ub_2_), K48-linked Ub_2_ ending with D77 on the C-terminus (K48-Ub_2_-D77), and corresponding monomeric ubiquitins. (**b**) MST binding analysis of substrates to PLpro^CoV-2^. Comparison of dissociation constants for the primary binding event (K_d1_). The K_d1_ is derived from a fit to three independent experiments with error bars corresponding to 68.3% confidence interval derived from error-surface projections. (**c**) Inter-and intramolecular non-specific cross-linking of PLpro^CoV-2^ complexes. Cross-linked samples with different molar ratios reveal formation of heterogeneous covalent PLpro^CoV-2^:Ub_2_ heterodimer complex bands (red, for glutaraldehyde cross-linker) compared to untreated reactions (black, control) by SDS-PAGE. (**d**) Quantification of the cleavage efficiency of hISG15 C-terminal fusions with peptides from Nsp2 (AYTRYVDNNF), Nsp3 (APTKVTFGDD), and Nsp4 (KIVNNWLKQL) from SARS-CoV-2 that mimic natural substrates of PLpro^CoV-2^, as revealed by SDS-PAGE gel. Shown is the percentage of the input population that has been cut by PLpro^CoV-2^. Experiment was performed in triplicate and is reported as an average with standard deviation. (**e**) Quantification of the cleavage efficiency of Nsp2 peptides as dependent on amino acid located at the C-terminus of “LRGG(X)” motif. Shown is the percentage of the input population that has been cut by PLpro^CoV-2^. Experiment was performed in triplicate and is reported as an average with standard deviation. (**f**) PLpro^CoV-2^ cleavage of K48-Ub_3_. SDS-PAGE gel reveals that PLpro cleaves Ub_3_ into Ub_2_ and Ub_1_ efficiently with various rates. (**g**) Mass spectrometry detection of cleavage patterns for PLpro^CoV-2^ hydrolyzing Ub_3_, in which the distal Ub (1) carries K48R mutation, the endo (2) Ub is ^15^N-labeled, and the proximal (3) Ub contains C-terminal D77 extension. Analysis of the time course reveals that this Ub_3_ is primarily hydrolyzed between Ubs 2 and 3. Masses of individual Ub units are shown on the top, and the identified products are shown at the bottom.

The earlier reported structure of the PLpro^CoV-1^:K48-Ub_2_ complex shows the proximal Ub bound to the Zn finger and palm domains via its surface hydrophobic patch (comprising residues L8, I44 and V70) placing the C-terminal tail modified with allylamine in a groove that is covalently linked to active site C111 (Supplementary Fig. 1d)^28^. A recent structure of full-length mISG15 bound to PLpro^CoV-2 27^ revealed a distinct binding mode of the proximal and distal UBL domains of mISG15. The proximal UBL is shifted away from the finger domain compared to the proximal-Ub binding mode while still placing the C-terminal LRGG tail into the active site of the protease (Supplementary Fig. 1d). Comparison of ISG15 and Ub recognition surfaces reveals that the hydrophobic patch centered on I44 in Ub (Supplementary Fig. 1b, c) is more polar in the proximal UBL of hISG15 and mISG15. Given that the prior studies used a triazole-linked Ub dimer we wanted to test the influence of the linker composition on binding to PLpro^CoV-2^. We used microscale thermophoresis (MST) binding experiments to quantify affinity between PLpro^CoV-2^ and three substrates: hISG15, K48-Ub_2_ and Ub_1_ (Fig. 1a). Additionally, we tested PLpro^CoV-2^ binding to K48-Ub_2_ and Ub_1_ containing a C-terminal aspartic acid (D77) after the “LRGG” PLpro recognition site (Fig. 1a), which is typically used for controlled enzymatic synthesis of ubiquitin chains^29^. Fitting our binding data to a 1:1 binding model (Supplementary Fig. 2a) resulted in abnormally high *χ*^2^ and systematic deviation in the residuals (Supplementary Fig. 2b). Improved fits were observed using a model that assumes two binding events with different binding constants (Supplementary Fig. 2a) yielding statistically significant reductions in the *χ*^2^ compared to one binding event for all datasets except Ub_1_-D77 (Supplementary Fig. 2b). We find that PLpro^CoV-2^ binds both hISG15 and K48-Ub_2_ with high affinity (90 nM and 70 nM, respectively) and more strongly than Ub_1_ (apparent K_d_, 170 nM) (Fig. 1b and Supplementary Table 1) although the actual microscopic K_d_ of Ub_1_ could be even lower as it may be able to bind to multiple sites on PLpro^CoV-2^ (see below). Interestingly, K48-Ub_2_ with a C-terminal aspartic acid (K48-Ub_2_-D77) binds almost tenfold weaker (600 nM) compared to Ub_2_, and Ub_1_-D77 exhibited weak binding (130 μM) (Fig. 1b and Supplementary Table 1). This is consistent with a lack of reported protease substrates with acidic residues at the C-terminus of the “LRGG(X)” motif^7^. Our analysis also suggests the presence of secondary binding events for hISG15 and K48-Ub_2_ with μM affinities (Supplementary Fig. 2a and Supplementary Table 1). This is supported by cross-linking data where we observed covalent adducts with molecular weight corresponding to heterodimers (PLpro:substrate) but also heterotrimers ((PLpro)_2_:substrate) (Fig. 1c and Supplementary Fig. 2c).

The observed heterogeneity of the species formed suggests that ISG15 may bind in a more defined orientation to PLpro while Ub_2_ appears to bind in several arrangements. As a comparison to Ub_1_, we also measured affinities for the isolated N-terminal (distal, hISG15_distal_) and C-terminal (proximal, hISG15_prox_) UBLs from hISG15 and found that they bind with micromolar dissociation constants similar to Ub_1_-D77 (Supplementary Fig. 2d and Supplementary Table 1) and consistent with NMR measurements (Supplementary Fig. 2e). The significantly weaker PLpro binding to the isolated UBLs compared to full-length hISG15 indicates that both UBL domains are required for the high-affinity binding of hISG15. These experiments indicate that there is a dominant binding mode between the substrate and PLpro but higher order complexes are also possible, thus justifying the need to fit our binding data with more complex binding models. This is more pronounced for Ub than for ISG15 as these different states appear to contribute to higher affinity of PLpro for Ub_2_ as compared to Ub_1_. This may be explained by the flexibility of the Ub-Ub (isopeptide) linker in K48-Ub_2_ that enables this dimer to adopt heterogeneous conformational ensembles^30–34^.

Derived from published structures of mISG15:PLpro^CoV-2^ and Ub_2_:PLpro^CoV-1 26–28^, we anticipated that hISG15 and Ub_2_ bind PLpro utilizing both UBL/Ub domains. However, the relatively small difference in Ub_2_ and Ub_1_ affinities suggests that the second Ub contributes modestly to the binding. Additionally, Ub_1_-D77 binds nearly three orders of magnitude more weakly compared to WT Ub_1_ while Ub_2_-D77 yields an affinity more similar to Ub_1_, perhaps indicative of a change in binding mode primarily utilizing a single Ub. To explore how affinity relates to PLpro proteolytic activity, we conducted PLpro^CoV-2^ cleavage assays for hISG15 with modified C-terminal tails mimicking natural SARS-CoV-2 substrates or K48-linked Ub_3_/Ub_2_. We found that PLpro^CoV-2^ can efficiently cleave peptides containing the “LRGG motif”. The hISG15 fusions to fragments of Nsp2, Nsp3 and Nsp4 are proteolyzed with similar rates (Fig. 1d and Supplementary Fig. 2f). We investigated how amino acid X (position 158) at the C-terminus of “LXGG↓(<u>**X**</u>)” motif can impact cleavage. When Ala of Nsp2 peptide is substituted with Glu, the fusion peptide is being cut the slowest (Fig. 1e and Supplementary Fig. 2g), consistent with binding measurements for Ub_1_-D77 and Ub_2_-D77 which have an additional acidic amino acid on C terminus (Fig. 1b and Supplementary Table 1). We also found that PLpro^CoV-2^ hydrolyzes K48-Ub_3_ to Ub_2_ and Ub_1_ rapidly, but the subsequent cleavage of Ub_2_ to Ub monomers is slow (Fig. 1f, g). Finally, we found that Ub_2_-D77 is cleaved more rapidly compared to Ub_2_, which suggests that despite Ub_2_-D77 binding with a notably lower affinity to PLpro^CoV-2^ it must be bound differently than Ub_2_ to enable the “productive” hydrolysis of Ub_2_ (Supplementary Fig. 2h). If Ub_3_ binds predominantly with two Ub units, there are two possible binding modes of Ub_3_ on PLpro.

To test which binding mode is dominant we used a K48-Ub_3_ that contains three “distinct” Ub units: 1 (mutant Ub-K48R, distal domain), 2 (U-^15^N-labeled Ub, endo (middle) domain) and 3 (Ub-D77, proximal domain) (Fig. 1g), allowing mass spectrometry-based identification of each cleavage product. Analysis by MS of a cleavage time course of this Ub_3_ construct reveals that the first cleavage occurs between Ub units 2 and 3, releasing the C-terminal Ub_1_-D77 (i.e. proximal) with only minor products for the other Ubs (Fig. 1g). This is consistent with Ub units 1 and 2 bound to PLpro with the C-terminal tail of unit 2 fitting the active site for rapid hydrolysis (Fig. 1g). Three Ub binding sites (S2, S1, and S1’) were proposed for PLpro^27^. In this model, Ub unit 1 would bind to S2, unit 2 to S1 and unit 3 to S1’ (Supplementary Fig. 1d). Interestingly, Ub_2_ could bind to PLpro in two modes. The high-affinity mode, as in Ub_3_ binding, where Ub unit 1 (distal) binds to S2 and unit 2 to S1, results in no cleavage. Only the second, lower-affinity mode is productive for Ub_2_ cleavage, wherein Ub unit 1 is bound to S1 and unit 2 (proximal Ub) located at S1’. In this arrangement the C-terminal tail of unit 1 connecting the two Ubs is placed in the active site of PLpro, and Ub_2_ is cut into monomers. These two modes of binding are competitive and because there is a difference in affinity, the rate of cleavage is reduced, but eventually resulting in complete disassembly of Ub_2_.

To further characterize the cleavage of Ub_3_ bound in different arrangements, we designed an NMR-based experiment using Ub_3_ or Ub_3_-D77 in which the endo Ub (unit 2) was ^15^N-labelled (as in the MS-based assay) allowing us to simultaneously monitor in real time the signal intensities for the isopeptide NH group of K48 (Supplementary Fig. 3a, red) and for G76 (Supplementary Fig. 3a, blue) of the endo Ub as a proxy for cleavage of Ub_3_ bound in two different geometries (Supplementary Fig. 3a, b). The signal corresponding to the conjugated C-terminal G76 of the endo Ub decreased rapidly (Supplementary Fig. 3c, blue triangles) and concomitantly with a rapid increase in its unconjugated G76 signal (Supplementary Fig. 3c, blue circles), indicating cleavage of the linker between the endo an proximal Ubs (units 2 and 3). By contrast, the K48 isopeptide signal disappeared much slower, indicating slower cleavage of the linker between the distal and endo Ub units (1 and 2), in full agreement with the MS-based cleavage assay (Fig. 1g, also 1f). For this sequence of Ub_3_ cleavage events to happen, Ub units 1, 2, and 3 must first occupy the S2, S1, and S1’ sites, respectively, to enable cleavage of the proximal Ub (unit 3), whereas the binding arrangement where Ub unit 1 occupies the S1 site and unit 2 the S1’ site, required for the distal-endo (1-2) linkage cleavage, occurs only after the proximal Ub (unit 3) gets cleaved.

Thus, our experiments uncover more complex alternate binding modes and sequence dependence of PLpro for two related substrates that have not been described to date. Our data also provide an alternative interpretation of recently published work^26,27^ as the composition and flexibility of the Ub-Ub linker can significantly impact the binding or cleavage or both for Ub_2_ and other protein substrates. This may explain the previously observed difference in affinity of PLpro^CoV-2^ between Ub_2_ and ISG15 which seems likely attributed to changes in mode of binding and/or conformational flexibility of the substrate linkage rather than mutations in the PLpro enzyme. Again, this is consistent with observed conformational flexibility of free K48-Ub_233,34_ and much lower conformational diversity of ISG15 as a free protein and in the complex with PLpro and other proteins (complex of hISG15 with the NS1 protein of influenza B virus^35^) (see below).

### Dual vs. single domain substrate recognition determines PLpro^CoV-2^ selectivity

We further investigated details of interaction between the PLpro^CoV-2^, hISG15 and K48-Ub_2_ to reveal similarities and differences. We determined crystal structures of PLpro^CoV-2^ with an active site C111S mutation, which inactivates PLpro, in complex with hISG15 at 2.98 Å resolution (Fig. 2a and Supplementary Table 2) and with K48-Ub_2_ at 1.88 Å resolution (Fig. 2b, Supplementary Fig 4a, and Supplementary Table 2). For the PLpro^CoV-2^:hISG15 complex we observe well resolved electron density for the proximal and distal UBL domains bound to S1 and S2 sites respectively (Fig. 2a). By contrast, for the PLpro^CoV-2^:K48-Ub_2_ structures determined at much higher resolution we observe strong electron density for the proximal Ub bound to S1 site with only weak signal for the distal Ub in S2 site and no electron density for Ub in S1’ site (Fig. 2b and Supplementary Fig. 4b). Despite weak electron density for the distal Ub, some regions of a density map resemble α-helix and amino acid chains corresponding to portions of the distal Ub from superposed structure of PLpro^CoV-1^ bound to K48-Ub_2_ (PDB id: 5E6J) upon only slight adjustment (Supplementary Fig. 4c). Therefore, there is sufficient room in our crystals of PLpro^CoV-2^-C111S:K48-Ub_2_ to fit the distal Ub at site S2 but the binding mode of the distal Ub may be less defined (Supplementary Fig. 4d)^28^. An alternative explanation is that PLpro^CoV-2^ (C111S) can hydrolyze the K48 linkage slowly, leading to a mixture of Ub_1_ and Ub_2_. To test this directly, we ran SDS-PAGE gels of our crystals and observed predominantly Ub_2_ with only a minor Ub_1_ species (Supplementary Fig. 4e).

**Figure 2.**
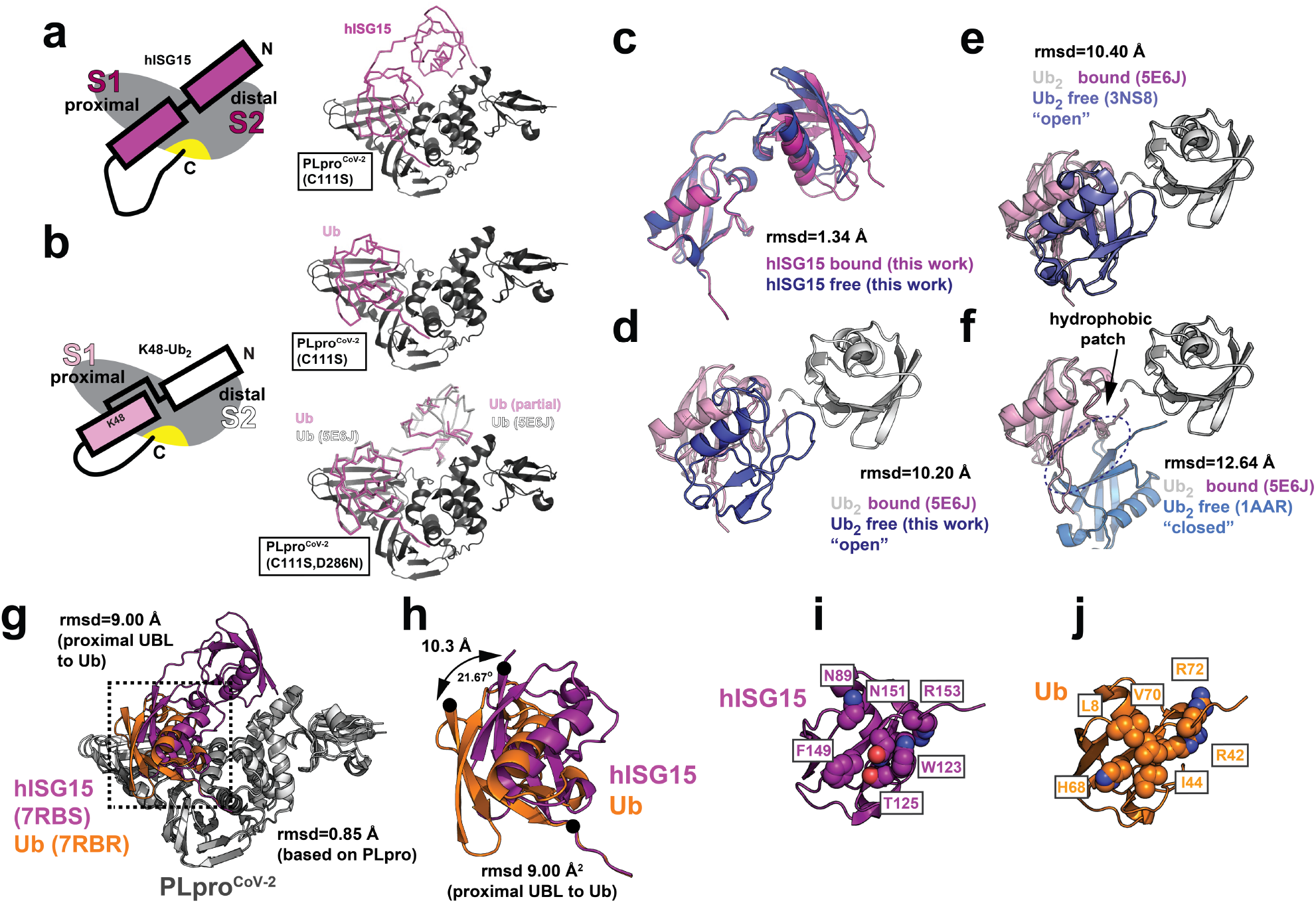
MX structure of PLpro^CoV-2^ bound to human ISG15 and K48-Ub_2_ reveals differential usage of distal domains. Schematic of PLpro bound to (**a**, left) hISG15 and (**b**, left) K48-linked Ub_2_. PLpro is shown in gray with an active site indicated in yellow. hISG15 is shown in magenta. Proximal and distal Ubs are pink and white, respectively. Crystal structures of PLpro^CoV-2^ bound to (a, right) hISG15 and (b, right) K48-Ub_2_. PLpro (C111S or C111S,D286N) is shown in cartoon representation, hISG15 and K48-Ub_2_ are shown as a backbone trace. PLpro, hISG15 and K48-Ub_2_ are colored as in (a, left) and (b, left). (**c**) Overlay of crystal structures of bound hISG15 (magenta, PDB id: 7RBS, this study) and unbound hISG15 (blue, PDB id: 7S6P, this study). (**d**) Overlay of bound conformation of K48-linked Ub_2_ observed in complex with PLpro^CoV-1^ (PDB id: 5E6J) with unbound conformation of K48-linked Ub_2_ (PDB id: 7S6O, this study). Proximal Ub of both bound and unbound conformations is shown in pink, distal Ub of bound conformation is shown in white, and distal Ub of unbound conformation is shown in blue. (**e**) Overlay of bound conformation of K48-linked Ub_2_ observed in complex with PLpro^CoV-1^ (PDB id: 5E6J) with unbound “open” conformation of Ub_2_ (PDB id: 3NS8). Ub units are shown as in (d). (**f**) Overlay of bound conformation of K48-linked Ub_2_ observed in complex with PLpro^CoV-1^ (PDB id: 5E6J) with unbound “closed” conformation of Ub_2_ (PDB id: 1AAR). Ub units are shown as in (d). Intramolecular Ub-Ub interface is indicated with an arrow. (**g**) Structural overlay of PLpro^CoV-2^:hISG15 and PLpro^CoV-2^:Ub_2_ (PDB id: 7RBR, this study). Proteins are shown in cartoon and colored gray (PLpro), magenta (hISG15) and orange (Ub_2_). (**h**) Zoom in of the boxed area in (g) representing overlay of the proximal domain of hISG15 and Ub. Proximal domain of hISG15 and Ub are represented as in (g). Rotation of the binding surface indicated with arrows. (**i**,**j**) Comparison of the binding surfaces of hISG15 (**i**) and Ub_1_ (**j**). Proximal domain of hISG15 and Ub are represented as in (g). Key residues are shown in space fill representation.

We have also determined structure of the PLpro^CoV-2^-C111S,D286N double-mutant:K48-Ub_2_ (Supplementary Fig. 4f and Supplementary Table 2). These crystals diffracted to higher resolution (1.45 Å) and allowed to model more fragments of distal Ub when compared to the 1.88 Å structure of PLpro^CoV-2^-C111S:K48-Ub_2_ (PDB id: 7RBR) (Supplementary Fig. 4f, g). In total 41 of 76 distal-Ub residues were built using this electron density map. Occupancies for most residues were lower and determined based on R-factor values and inspection of F_o_ − F_c_ difference maps. The location of the distal Ub fragment closely matches the position of distal Ub in the structure of PLpro^CoV-1^ bound to a K48-Ub_2_ (PDB id: 5E6J). Interestingly, including distal Ub from PLpro^CoV-1:^K48-Ub_2_ (PDB id: 5E6J) into structure lowers R-work factor by 0.32% with slight increase of R-free factor by 0.12% as compared for the structures with and without distal Ub. By contrast, the insertion of distal Ub fragments using the current structure resulted in decrease of both R-work/R-free factors by 0.64/0.11%, respectively. It is important to mention that insertion of complete distal Ub causes clashes with neighboring molecules, suggesting that multiple conformations must be present in the crystal of PLpro^CoV-2^-C111S,D286N:K48-Ub_2_. Our structures revealed how the protease differentially recognizes hISG15 using both UBL domains while K48-Ub_2_ is predominantly recognized using the proximal Ub.

We additionally determined structures of the free hISG15 and K48-Ub_2_ to 2.15 Å and 1.25 Å resolution, respectively. The bound and free hISG15 conformations are similar with a root-mean-square deviation (rmsd) for Cα atoms of 1.34 Å (Fig. 2c) based on alignment of the C-terminal (proximal) UBL. An analogous comparison of our unbound K48-Ub_2_ structure with the only known PLpro-bound conformation of K48-Ub_2_ (from SARS-CoV-1)^28^ revealed an rmsd of 10.2 Å. The Ub units are oriented very differently relative to each other (Fig. 2d), with the PLpro-bound Ub_2_ in an extended conformation (like hISG15) while the unbound Ub_2_ in an “open” conformation with the functional hydrophobic surface patches exposed for binding. We also compared the “bound” conformation with two other published canonical “open” (Fig. 2e) and “closed” (Fig. 2f) conformations of K48-Ub_2_ revealing large differences in rmsd of 10.4 Å (PDB id: 3NS8) and 12.64 Å (PDB id: 1AAR), respectively. In the “open” conformation the functional binding surface on Ub is exposed while in the “closed” conformation the functional nonpolar surfaces are engaged in intramolecular Ub-Ub interactions (Fig. 2f, arrow). Interestingly, our new K48-Ub_2_ structure is nearly identical to the previously reported “open” conformation (PDB id: 3NS8) with an rmsd of 0.23 Å^30^. These data reflect that hISG15 is more rigid while the K48-Ub_2_ exists as ensemble of conformational states which may influence binding and recognition by PLpro and other USPs^31–33^. Moreover, different Ub linkage types may exploit different interdomain conformational space explaining the source of specificity^34^. These observations confirm that PLpro is capable of recognizing distinct surfaces presented on Ub or UBL dimers (see below).

### Functional surfaces of hISG15 and K48-Ub_2_ are recognized differentially by PLpro^CoV-2^

We first compared the binding modes of hISG15 and proximal Ub in our structures. The structures of PLpro in both complexes are similar with an rmsd of 0.85 Å, but the binding surface contacts of the proximal Ub are shifted towards the fingers domain of PLpro compared to the proximal UBL domain of ISG15 (Fig. 2g, h). This shift in the binding mode is manifested by a 21.7° rotation around the C-termini of hISG15 and Ub displacing the N-terminal residue by 10.3 Å (Fig. 2h). This is despite the structural homology between the hISG15 proximal UBL domain and Ub (rmsd of 0.96 Å) (Supplementary Fig. 5a); thus it is likely dictated by differences in PLpro binding to the Ub and UBL interacting surfaces (Supplementary Fig. 5b).

We also compared our structure to the previously published PLpro^CoV-2^:mISG15^26^ complex. Overlay of the two structures (PDB ids: 7RBS and 6YVA) reveals good structural similarity with the overall rmsd of 0.70 Å for PLpro and 1.40 Å for hISG15 and mISG15 (Supplementary Fig. 6a). The proximal UBL domains of both ISG15s are well aligned and make several conserved interactions with the PLpro but interaction with the distal UBL domain shows the largest deviation (Supplementary Fig. 6a). We compared the contacts from the distal domain of mISG15 and hISG15 to previously determined hotspot residues (F69 and V66) on PLpro^CoV-2 26^. We find that in the PLpro^CoV-2^:mISG15 structure K30 and M23 of the distal UBL of mISG15 interact with F69 of PLpro^CoV-2^, while V66 of PLpro^CoV-2^ interacts with A2 of the substrate (Supplementary Fig. 6b). By contrast, in our new PLpro^CoV-2^:hISG15 structure residue 30 of hISG15 is an alanine, thus leaving M23 alone to stabilize the interaction with F69 of PLpro^CoV-2^, while the N-terminus of hISG15 interacts with V66 of PLpro^CoV-2^ (Supplementary Fig. 6c). Residue 20 in ISG15 makes similar nonpolar contacts with V66 but it varies between the mouse (T20) and human (S20) protein (Supplementary Fig. 6b, c). We additionally compared the interactions between the proximal UBL domains of the mISG15 and hISG15 where the UBL binds in a similar binding mode (Supplementary Fig. 6d). We find that overall, the two ISG15 proteins make similar, but not identical contacts determined by the sequence variation between mISG15 and hISG15. This suggests that the virus may have a different impact if it infects distinctive species. The central interacting residues on PLpro^CoV-2^ are Y171, E167 and M208 which interact with conserved R153/R151, W123/W121 and P130/P128 on the proximal domains of hISG15 and mISG15, respectively (Supplementary Fig. 6e, f). The interaction is centered on a salt bridge between E167 of PLpro^CoV-2^ and R153 of hISG15 while the equivalent arginine (R151) in mISG15 is not oriented properly to form a salt bridge. Nonetheless, this core interaction is stabilized by nonpolar interactions of the surrounding residues from both sides of the interface including Y171 of PLpro^CoV-2^ and W123/W121 and P130/P128 from hISG15 and mISG15, respectively (Supplementary Fig. 6e, f). By contrast, interactions with R166 of PLpro^CoV-2^ vary more significantly between mISG15 and hISG15. In the mISG15 structure the side chain of M208 is not resolved while in the hISG15 structure M208 packs against R166 (Supplementary Fig. 6f).

Interestingly, R166 forms a salt bridge with E87 of mISG15 which is changed to asparagine (N89) in hISG15 (Supplementary Fig. 6e, f). To compensate for this loss of interaction, N151 of hISG15 makes a hydrogen bond with R166 (Supplementary Fig. 6e, f). This highlights subtle sequence changes between mISG15 and hISG15 that allow interface rearrangements while preserving the binding mode and may explain reported previously differences in binding between human and mouse ISG15^26^. Our structures show that hISG15 binds PLpro^CoV-2^ utilizing both proximal and distal UBL domains (Fig. 2a), while binding of K48-Ub_2_ is primarily driven by interaction with the proximal Ub with only weak density observed for the distal Ub (Fig. 2b and Supplementary Fig. 4b). We also compared our PLpro^CoV-2^:hISG15 structure to a recent structure of PLpro^CoV-2^ bound to only the proximal domain of hISG15^27^ (Supplementary Fig. 7a; PDB id: 6XA9, 2.9 Å resolution). As in the PLpro^CoV-2^:mISG15 complex, the structural similarity is high, with an overall Cα rmsd of 1.0 Å. A comparison of the interface contacts reveals nearly identical interactions, even preserving side-chain rotamers between the proximal hISG15 and PLpro in the two structures (Supplementary Fig. 7b). Finally, our new structure of PLpro^CoV-2^:K48-Ub_2_ is nearly identical in binding mode to the previously published structure of PLpro^CoV-2^:Ub_1_ (Supplementary Fig. 7c; PDB id: 6XAA, 2.7 Å resolution) with a Cα rmsd of 0.32 Å and nearly identical side chain rotamers at the interface (Supplementary Fig. 7d). Interestingly, our structure was determined to higher resolution and without the introduction of a covalent linkage of Ub_2_ to PLpro^CoV-2^ suggesting that the covalent linkage does not alter the physiological binding of the substrate. By contrast, however, introduction of a synthetic Ub-Ub linker in Ub_2_ does influence binding of the substrate to PLpro^CoV-2^.

However, PLpro cuts K48-Ub_2_ slowly, because for productive cleavage the distal Ub must bind to S1 (low affinity binding) and proximal Ub occupy the S1’ site. S1’ site is likely less specific as it must accept multiple protein sequences. This is supported by our observations discussed earlier that K48-Ub_3_ is cut efficiently to Ub_2_ and Ub, with the remaining Ub_2_ bound in a non-cleavable binding mode.

### Interactions of K48-Ub_2_ and ISG15 with PLpro^CoV-2^ in solution using NMR

We then used NMR to further characterize PLpro^CoV-2^ binding to hISG15 and K48-Ub_2_ and to examine if the contacts observed in crystals also occur in solution. The addition of unlabeled PLpro^CoV-2^-C111S caused substantial perturbations in the NMR spectra of ^15^N-labeled hISG15 (Fig. 3a). We observed disappearance of signals of free hISG15 and emergence of new ones; this indicates slow-exchange binding regime^36,37^, consistent with the sub-μM K_d_ values measured by MST (Fig. 1b and Supplementary Table 1). The strongly attenuated signals of hISG15 residues, including the C-terminal G157, are consistent with our crystal structure of the PLpro^CoV-2^:hISG15 complex (Fig. 3a, h). A similar behavior was observed for ^15^N-labeled K48-Ub_2_ upon addition of PLpro^CoV-2^ (Fig. 3b, c), where both the distal and proximal Ubs exhibited strong attenuation or disappearance of NMR signals and emergence of new signals, primarily for residues in and around the hydrophobic patch as well as the C-termini. The affected residues mapped to the binding interface in our PLpro^CoV-2^:K48-Ub_2_ crystal structure (Fig. 3i), and the slow-exchange behavior is also consistent with the sub-μM K_d_ values (Fig. 1b and Supplementary Table 1). The slow-exchange binding regime for both hISG15 and K48-Ub_2_ is also generally consistent with the reported slow off-rates (0.2, 0.4 s^-1^)^26^. PLpro^CoV-2^ also caused noticeable perturbations in the NMR spectra of Ub_1_, although these were weaker than in Ub_2_, and several residues showed gradual signal shifts indicative of fast exchange, consistent with our MST measurements (Fig. 3d, Fig. 1b, and Supplementary Table 1). We additionally compared binding between PLpro^CoV-2^ and ^15^N-labeled hISG15_distal_ and hISG15_prox_ and detected few changes in the spectra of hISG15_distal_ indicative of weak interactions (Supplementary Fig. 2e). Interestingly, titration of PLpro^CoV-2^ into ^15^N-labeled hISG15_prox_ yielded similar signal shifts and attenuations to those observed in full-length hISG15, suggesting similar binding interactions, although unbound signals were still observed even at 2x molar access of PLpro, consistent with weaker affinity. Notably, the signal attenuation of C-terminal G157 and the characteristic shifts of PLpro^CoV-2^ Trp signals indicate that the C-terminus of hISG15_prox_ alone can still bind to the active site of PLpro^CoV-2^ (Supplementary Fig. 2e), consistent with a slightly higher affinity of PLpro^CoV-2^ for hISG15_prox_ compared to hISG15_distal_ (Supplementary Fig. 2d).

**Figure 3.**
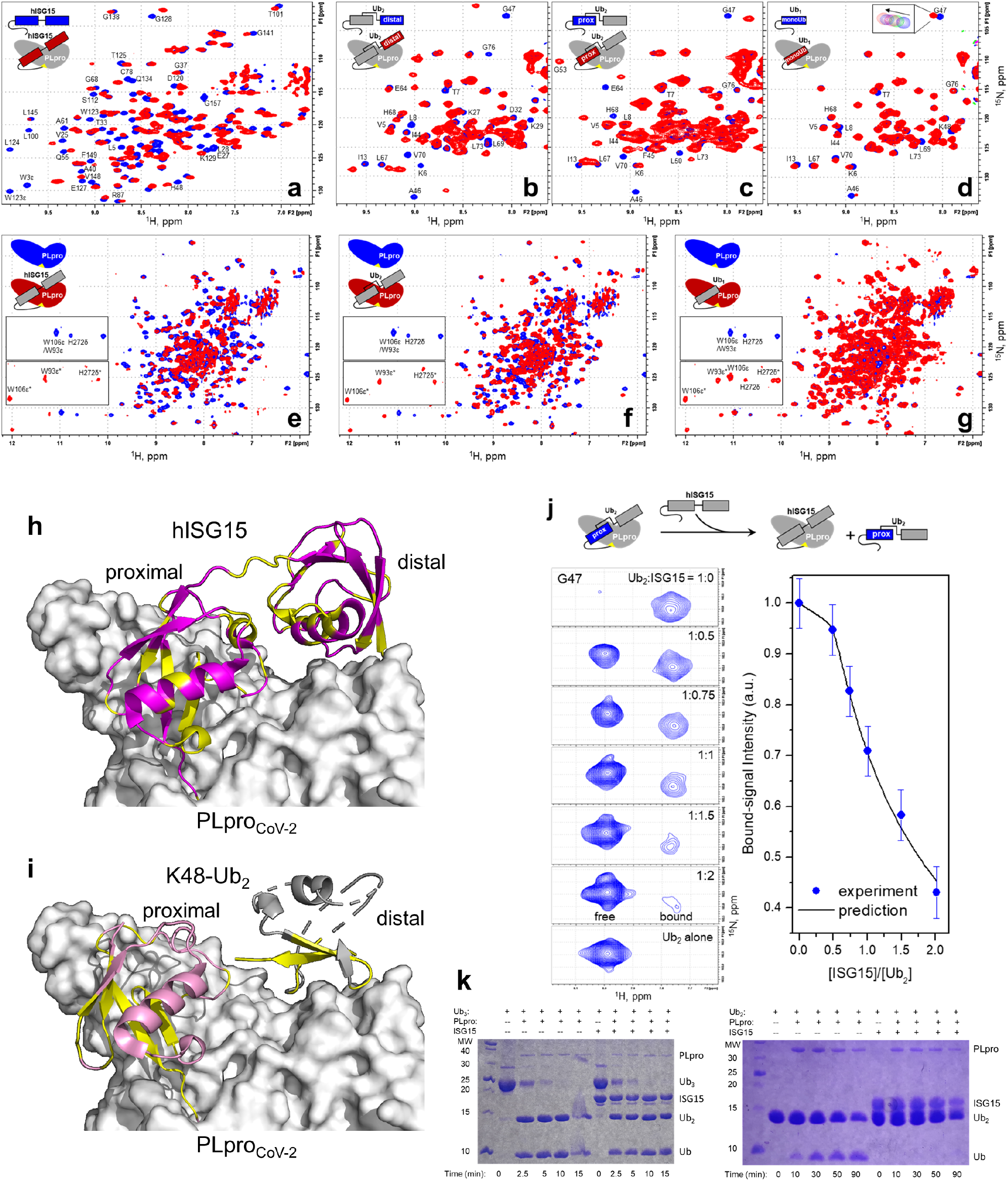
NMR data showing PLpro^CoV-2^ interactions with hISG15 and K48-Ub_2_. (**a-d**) Overlay of ^1^H-^15^N SOFAST-HMQC spectra of ^15^N-labeled (**a**) hISG15, (**b**) distal Ub in Ub_2_, (**c**) proximal Ub in Ub_2_, and (**d**) monomeric Ub, alone (blue) and with 1.2 molar equivalents of unlabeled PLpro^CoV-2^ (C111S) (red). Signals of select residues are indicated. The inset in d illustrates gradual shift of G47 signal during titration. **(e-g)** Overlay of ^1^H-^15^N TROSY spectra of ^15^N-PLpro^CoV-2^ (C111S,Y171H) alone (blue) and (in red) with 1.5 molar equivalents of unlabeled (**e**) hISG15, (**f**) Ub_2_, and (**g**) monomeric Ub (red). Insets zoom on the region containing indole HN signals of tryptophans (W93 and W106) and HN of imidazole ring of histidine H272. (**i**, top panel) hISG15 residues with strong signal perturbations mapped (yellow) on our structure of PLpro^CoV-2^:hISG15 complex. (**i**, bottom panel) Residues in the proximal and distal Ubs with strong signal perturbations mapped (yellow) on our structure of PLpro^CoV-2^:K48-Ub_2_ complex. (**j**) Competition for PLpro^CoV-2^ binding between Ub_2_ and ISG15. Shown on the left are representative ^1^H-^15^N NMR signals of G47 in ^15^N-labeled proximal Ub of Ub_2_ mixed with unlabeled PLpro^CoV-2^ (C111S,Y171H) (in 1:1.5 molar ratio) upon addition of unlabeled hISG15, for indicated values of hISG15:Ub_2_ molar ratio. Shown on the right is the intensity of the PLpro-bound signal of G47 as a function of [hISG15]:[Ub_2_] (dots) and the predicted molar fraction of bound Ub_2_ (line) using the K_d1_ values obtained in this work (Supplementary Table 1). (**k**) SDS-PAGE gels showing the inhibitory effect of hISG15 on disassembly of K48-linked Ub_2_ by PLpro^CoV-2^ (right gel) and minimal effect (if any) of hISG15 on disassembly of K48-linked Ub_3_ (left gel). The hISG15 and Ub_2_ constructs used here all had G (G157 or G76) as the C-terminal residue.

We also performed reverse-titration NMR experiments where unlabeled hISG15, K48-Ub_2_, or Ub_1_ was added to ^15^N-labeled PLpro^CoV-2^. Both hISG15 and K48-Ub_2_ caused substantial perturbations in the ^15^N-PLpro^CoV-2^ spectra (Fig. 3e, f). Particularly noticeable was the change in the indole NH signals of W93 and W106 located in close proximity to the active site of PLpro, as well as of imidazole NH signal attributed to the active site H272 (Fig. 3e, f, Supplementary Fig. 8a, b), in agreement with the C-termini of hISG15 and Ub_2_ entering the active site of PLpro in our crystal structures. The addition of Ub_1_ caused significantly lesser overall ^15^N-PLpro^CoV-2^ signal perturbations, although the W_ε_ and H_δ_ signal shifts were clearly visible when Ub_1_ was in significant excess (Fig. 3g, Supplementary Fig. 8c-e). Even at 8-molar excess of Ub_1_ both free and bound W_ε_/H_δ_ signals were present, consistent with weaker binding. Taken together, the NMR data qualitatively suggest that the apparent strength of PLpro^CoV-2^ binding is: hISG15 ≈ K48-Ub_2_ > Ub_1_, consistent with our MST data.

These NMR data indicate that binding to PLpro involves both UBLs of hISG15 and both Ubs of K48-Ub_2_. Interestingly, despite being identical and having very similar chemical shifts in the unbound state (Supplementary Fig. 9a), the distal and proximal Ubs show markedly different signal perturbations indicative of distinct contacts with PLpro (Fig. 3b, c, and Supplementary Fig. 9b). While the perturbed residues in the proximal Ub agree well with our PLpro^CoV-2^:K48-Ub_2_ crystal structure, where this Ub occupies the S1 site, several perturbations observed in the distal Ub (most notably for N25, K27, K29, D32, K33 in the α-helix) are not observed in the PLpro^CoV-2^:K48-Ub_2_ or PLpro^CoV-1^:K48-Ub_2_ crystal structures. Thus, we cannot exclude possible additional modes of interaction between the distal Ub and PLpro^CoV-2^. This might explain the low electron density for the distal Ub in the PLpro^CoV-2^:K48-Ub_2_ crystal structure. Interestingly, for several residues in Ub_1_ the shifted signals upon addition of PLpro^CoV-2^ appear at positions intermediate between those in the distal and proximal Ubs of K48-Ub_2_ (Supplementary Fig. 9b), suggesting that Ub_1_ might be sampling both S1 and S2 sites on PLpro.

In agreement with the MST results, our NMR data demonstrate that placement of an aspartate at the C-terminus of hISG15, K48-Ub_2_, and Ub_1_ reduced substantially their affinity for PLpro^CoV-2^, as evident from noticeably weaker NMR signal perturbations observed in both the D-extended substrates and PLpro (Supplementary Fig. 10). It should be mentioned that in all the NMR studies presented here the addition of PLpro^CoV-2^ resulted in the overall NMR signal broadening/attenuation reflecting an increase in the size (hence slower molecular tumbling) upon complexation with a ∼36 kDa protein. The finding that hISG15 and K48-Ub_2_ bind to the same sites on PLpro^CoV-2^ enabled us to directly compare their affinities for PLpro^CoV-2^ in a competition assay where hISG15 was added to a preformed PLpro^CoV-2^:K48-Ub_2_ complex, and the bound state of Ub_2_ was monitored by ^1^H-^15^N NMR signals of ^15^N-labeled proximal Ub (Fig. 3j). Titration of unlabeled hISG15 into a 1:1.5 mixture of K48-Ub_2_ and PLpro^CoV-2^ resulted in gradual disappearance of PLpro-bound signals of Ub and concomitant emergence of free K48-Ub_2_ signals at their unbound positions in the spectra (Fig. 3j, Supplementary Fig. 9c). The observed decrease in the intensity of the bound signals agrees with the prediction based on the K_d1_ values for K48-Ub_2_ and hISG15 derived from our MST experiments (Fig. 3j) but not with the K_d_ values reported previously^26^ (Supplementary Fig. 9d).

Since hISG15 binds to the same PLpro^CoV-2^ surface as Ub_2_ but contains an uncleavable linkage between UBLs, we then examined if hISG15 can inhibit polyUb cleavage by PLpro. When hISG15 was added to a cleavage reaction of Ub_3_ or Ub_2_ by PLpro^CoV-2^ it did not interfere with Ub_3_ cleavage to Ub_2_ (Fig. 3k, left gel) but it blocked hydrolysis of Ub_2_ to monomers (Fig. 3k, right gel, also Supplementary Fig. 9e). A similar effect was observed on cleavage of Ub_2_ in the presence of Ub_1_ (Supplementary Fig. 9e). This can be explained by different options for productive cleavage of Ub_3_ and Ub_2_. The productive cleavage of Ub_3_ is predominantly accomplished by binding of two Ub units (2 and 1) to S1 and S2 sites on PLpro^CoV-2^, respectively, and the unit 3 of Ub_3_ occupying the S1’ site, thus placing the isopeptide bond on its K48 in the active site of PLpro^CoV-2^. This is a high affinity Ub_3_ binding, and hISG15 and particularly Ub_1_ cannot easily compete for binding. As discussed earlier, K48-Ub_2_ can bind in two different modes, one with two Ub domains binding to S1 and S2 sites on PLpro, but this binding cannot result in cleavage. In order to break the isopeptide bond between two Ubs the distal Ub must bind to the S1 site on PLpro such that the proximal Ub will then occupy the S1’ site. The LRGG motif can then be recognized, and the isopeptide bond is cleaved. But Ub_2_ binding through a single Ub unit to S1 site is of low affinity, thus both hISG15 and Ub_1_ can compete with Ub_2_ and inhibit its cleavage.

### Specific contacts in PLpro:substrate complexes detected with XL-MS

Our structural experiments indicate differences in how hISG15 and Ub_2_ are recognized by PLpro^CoV-2^. To gain more insight into the proposed dynamics of the interactions, we employed a XL-MS approach (Fig. 4a). We found that the 4-(4,6 dimethoxy-1,3,5-triazin-2-yl)-4-methyl-morpholinium chloride (DMTMM) cross-linker produced robust heterodimers of PLpro^CoV-2^ with hISG15, K48-Ub_2_, Ub_1_ or K48-Ub_2_-D77 (Fig. 4b and Supplementary Fig. 11a). We identified two contacts between D61 and D62 on PLpro^CoV-2^ to K35 on the distal UBL of hISG15 (Fig. 4c, 19 and 43 contacts) which map well onto our structure with the distances between carboxylates (D61 and D62) and N_ζ_ (K35) of 14.9 and 8.6 Å, respectively (Fig. 4c) with C_β_-C_β_ distances below 30 Å consistent with the cross-linker geometry^38^. By contrast, we detected 19 cross-links between PLpro and K48-Ub_2_, however, due to the sequence degeneracy between the two Ubs we interpreted the data based on shortest distance (Supplementary Fig. 11b). Using this strategy, 12 of the 19 observed contacts fall below a 30 Å threshold (Fig. 4d). Of these 12 contacts, 7 involve K6 from the distal Ub to the N-terminal thumb domain of PLpro^CoV-2^, including E70, consistent with the distal Ub binding mode seen in the PLpro^CoV-1^:Ub_2_ structure (PDB id 5EDJ)^28^. Additionally, of the 12 contacts, two between K190 on the fingers domain of PLpro^CoV-2^ and E64 and E18 of Ub have 16 Å and 28.6 Å Cβ-Cβ distances which are compatible with the placement of the proximal Ub in the S1 binding site (Fig. 4d). Similarly to what NMR indicated, for Ub_1_ we found 23 cross-links that localize to both S1 and S2 sites (Supplementary Fig. 11c). To find alternate binding sites that explain 7 of 19 identified cross-links from the PLpro^CoV-2^:K48-Ub_2_ dataset, we modeled 5,000 binding modes of the distal Ub by docking a Ub monomer to PLpro:Ub_prox_ complex containing (proximal) Ub in the S1 binding site and utilizing constraints that place the docked (distal) Ub with its C-terminus in proximity to K48 of the proximal Ub (Fig. 4e, g, blue spheres). We compared the energy of the complex as a function of the sum of distances between 7 cross-linked atoms pairs that were unexplained in the initial model (Supplementary Fig. 11d). A low-energy model that explains 4 of the 7 cross-links (Fig. 4f) localizes the Ub to the PLpro’s UBL and thumb domains, a site that we named S2’ (Fig. 4g). In a parallel docking approach, we assumed an alternative binding mode where the distal Ub was placed in the S1 binding site (Supplementary Fig. 11e), and applied a constraint from the C-terminus of that distal Ub to K48 of the docked (proximal) Ub. Comparison of the energetics and sum of cross-linked distances, uncovered a low-energy model with the docked Ub near the PLpro’s UBL that similarly explains 4 of 7 contacts (Supplementary Fig. 11e, f).

**Figure 4.**
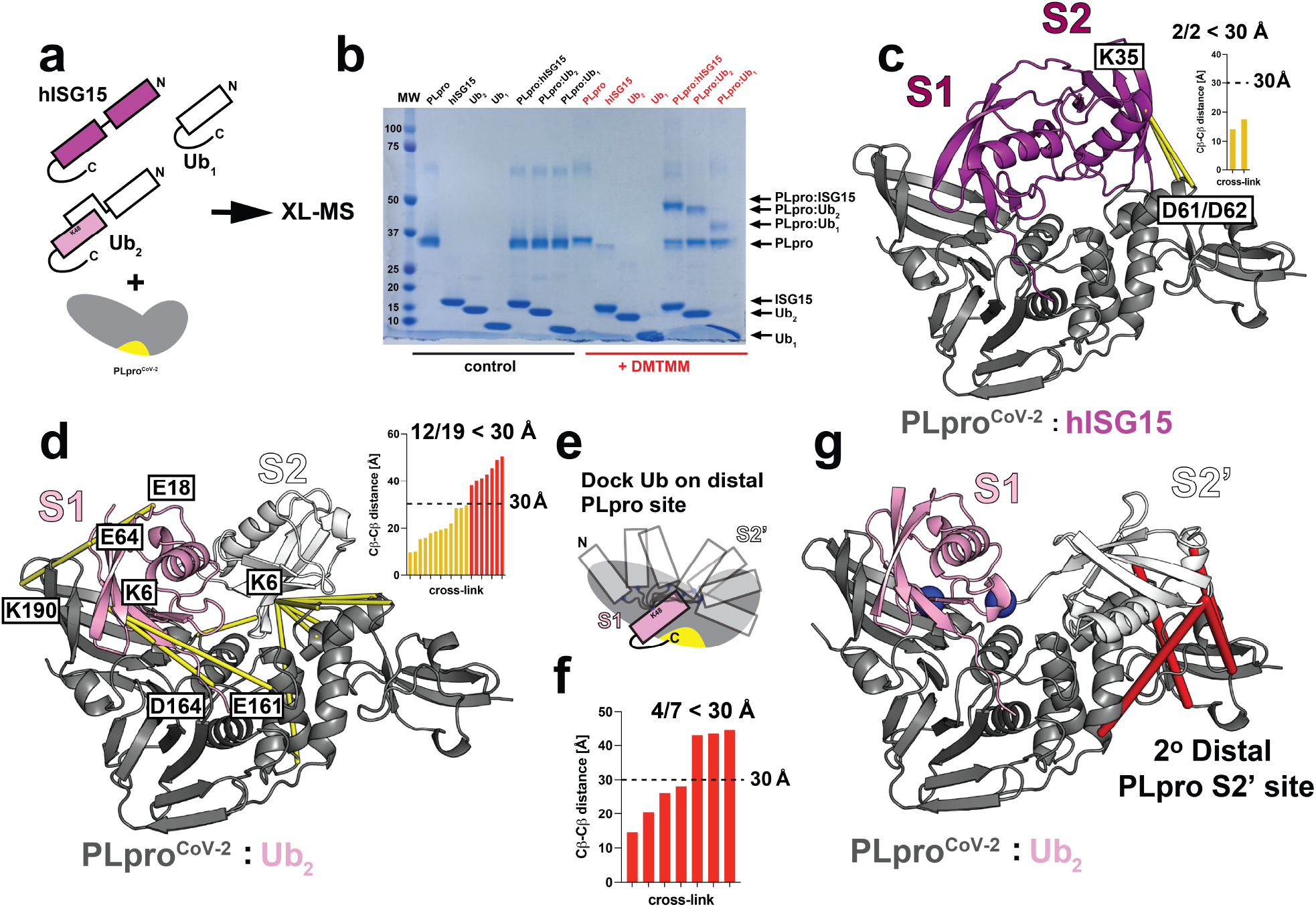
Cross-linking mass spectrometry (XL-MS) analysis of PLpro in complex with hISG15 and K48-Ub_2_. (**a**) Schematic illustration of cross-linking mass spectrometry experiments for heterodimer complexes of PLpro^CoV-2^ with hISG15, K48-Ub_2_, and Ub_1_. Active site of PLpro^CoV-2^ is indicated in yellow. (**b**) Cross-linked samples reveal formation of covalent heterodimer complex bands (red, DMTMM cross-linker) compared to untreated reactions (black, control) by SDS-PAGE. (**c**) Interactions between PLpro^CoV-2^ (D61/D62) and the distal hISG15 domain (K35) identified by XL-MS mapped on the structure of heterocomplex. Both identified contacts are shorter than 30 Å. PLpro (gray) and hISG15 (magenta) are shown in cartoon representation. Cross-links are colored yellow. (**d**) Interactions between PLpro^CoV-2^ and K48-Ub_2_ identified by XL-MS mapped on the structure of the heterocomplex. Twelve contacts found to be shorter than 30 Å show interaction of K48-Ub_2_ with fingers, palm and thumb domains of PLpro^CoV-2^. PLpro^CoV-2^ is shown in cartoon representation and colored gray. K48-Ub_2_ is shown in cartoon representation and colored white and pink for the proximal and distal Ubs, respectively. Cross-links are colored yellow. (**e**) Modeling strategy showing docked Ub monomer to PLpro^CoV-2^:Ub complex with Ub placed into the S1 binding site. Utilized constraint maintaining proximity between K48 of the proximal Ub and C-terminus of the docked Ub is shown in blue spheres. (**f**) A low energy model generated with strategy presented in (e) explains 4 of the remaining 7 contacts from the XL-MS in (d). (**g**) Contacts shorter than 30 Å between low energy model of PLpro^CoV-2^ bound to Ub_2_ in the S1 and S2’ sites as shown in (e, f). Cross-links between distal Ub and thumb and UBL domains of PLpro^CoV-2^ are colored red. Constraint used in docking (e) is indicated as blue spheres. PLpro^CoV-2^ and K48-Ub_2_ are shown as in (d).

To resolve this discrepancy in binding modes, we performed XL-MS on PLpro^CoV-2^:Ub_2_ in which only the proximal Ub was uniformly ^15^N labeled. Analysis of cross-links containing the unlabeled (distal) Ub uncovered 14 cross-links (Supplementary Fig. 11g) of which 8 are compatible with placement of this Ub in the S2 site and 7 of these again involve K6 interacting with the N-terminal thumb domain similar to the data collected with unlabeled Ub_2_ (Fig. 4d). This allowed us to propose that K48-Ub_2_ can bind to PLpro in two different binding modes (Fig. 4e and Supplementary Fig. 11e). Having more confidence that the distal Ub is bound in the S2 site, we again mapped the remaining 6 unexplained contacts from proximally ^15^N labeled dataset onto the docked model in which we sampled movement of the distal Ub (Fig. 4e); this model can explain 4 additional contacts (Supplementary Fig. 11h). Finally, we also interpreted XL-MS data for PLpro^CoV-2^:Ub_2_-D77 and using a model derived from the sampling of the S1’ site (Supplementary Fig. 11e). We can explain the two contacts consistent with alternate binding modes of Ub_2_-D77 (Supplementary Fig. 11i) with one contact comprising the distal Ub bound to the S1 site and the second involving the proximal Ub bound to the UBL domain of PLpro (S1’ site) to produce a cleavage-competent binding mode. This change in binding mode may explain faster cleavage kinetics of Ub_2_-D77 compared to Ub_2_ (Supplementary Fig. 2h). Our combined experiments not only reaffirm the dominant binding modes between PLpro and hISG15 and K48-Ub_2_ (to S1 and S2 sites) but also begin to clarify alternate binding modes that explain the heterogeneity of binding to S1 and S1’ sites.

### Identification of specificity-determining sites in PLpro^CoV-2^:Ub_2_ and PLpro^CoV-2^:hISG15 complexes

To better understand the energetic contribution of the residues at the PLpro:substrate interfaces, we applied an *in silico* alanine scan approach^39^ to PLpro^CoV-2^ in complexes with hISG15 and K48-Ub_2_ (Fig. 5a). We first identified PLpro interface residues that contact the substrates and employed Rosetta^39^ to calculate ΔΔG_binding_ comparing WT and alanine mutants. Analysis of ΔΔG_binding_ for PLpro^CoV-2^ with the two substrates revealed interaction hotspots, most notably PLpro^CoV-2^ residues E167/R166/Y264 and F69, for stabilizing S1 and S2 Ub/UBL binding sites in K48-Ub_2_ and hISG15 substrates (Fig. 5b, c). Additionally, we find that for PLpro^CoV-2^, the S2 site has a preference for hISG15 (Fig. 5b, c, colored in red) but the S1 site has an overall preference for Ub (Fig. 5b, c, colored in blue). We additionally, interpreted the interface energetics for models of PLpro^CoV-2^ in complex with Ub_2_ using two geometries of the distal Ub derived from the PLpro^CoV-1^:Ub_2_ and our PLpro^CoV-2^-C111S,D286N:Ub_2_ with a partial distal Ub (Supplementary Fig. 12, described in Methods). For K48-Ub_2_ the primary interaction is with the proximal Ub domain in S1 site but additional interaction comes also from the distal Ub interacting with S2 site contributing to stronger binding. In protein complexes, residues that surround protein interaction hotspots typically play important roles in determining specificity^40^. Indeed, in the S1 site, Y171 provides more stabilization for hISG15 compared to Ub_2_ (Fig. 5b). These analyses highlight how prediction of binding energetics combined with structural data can help interpret dynamics of domain binding.

**Figure 5.**
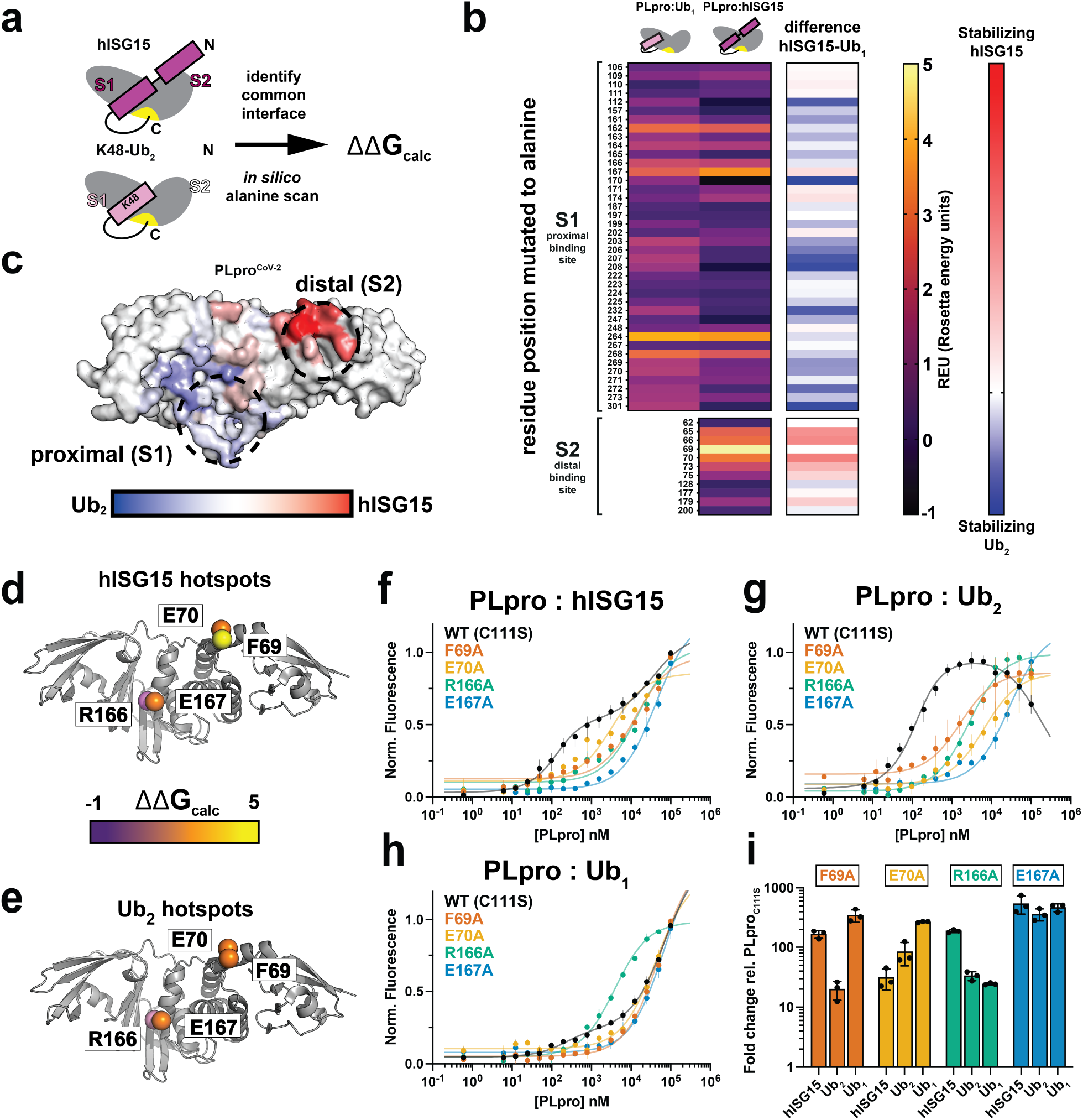
Prediction and validation of specificity determining surfaces on PLpro^CoV-2^. (**a**) Schematic illustration of identification of substrate binding surfaces on PLpro^CoV-2^ and *in silico* mutagenesis for heterodimer complexes with hISG15 and K48-Ub_2_. (**b**) Heat-map results of ΔΔG_binding_ calculations of *in silico* alanine scan for PLpro^CoV-2^ in complex with hISG15 or Ub_1_. Interface residue positions in the S1 (proximal) and S2 (distal) binding sites are labeled. The heat-map is colored from black to yellow. The last column represents results of calculations of difference between REU (Rosetta energy units) for PLpro^CoV-2^:hISG15 and PLpro^CoV-2^:K48-Ub_2_ and is colored from blue to red. (**c**) Results of *in silico* mutagenesis for complexes of PLpro^CoV-2^ with hISG15 and K48-Ub_2_ mapped on surface representation of PLpro^CoV-2^. Hotspot sites identified as those driving stability towards hISG15 are colored red, and those driving stability towards K48-Ub_2_ are colored blue. Summary of PLpro^CoV-2^ alanine mutants (F69A, E70A. R166A and E167A) tested for binding to hISG15 (**d**), Ub_2_ (**e**) and Ub_1_. Mutants are shown as Cα spheres and are colored according to ΔΔG_calc_ from black to yellow. (**f-h**) PLpro^CoV-2^ WT (C111S) (black), PLpro^CoV-2^ F69A (orange), PLpro^CoV-2^ E70A (yellow), PLpro^CoV-2^ R166A (green) and PLpro^CoV-2^ E167A (blue) titrations with hISG15 (f), Ub_2_ (g) and Ub_1_ (h). Data is shown as triplicates and is plotted as the average with the range of individual replicates. Data is fitted to the preferred 1:2 binding model for WT (C111S) and to 1:1 binding model for mutants using PALMIST. (**i**) Summary of fold change in K_d_s calculated as a ratio between PLpro^CoV-2^ WT (C111S) and PLpro^CoV-2^ F69A, PLpro^CoV-2^ E70A, PLpro^CoV-2^ R166A, PLpro^CoV-2^ E167A to hISG15, Ub_2_ and Ub_1_. Data is shown as triplicates and is plotted as averages with standard deviation.

Our *in silico* analysis of the PLpro:substrate interfaces predicts hotspot sites that are important for PLpro^CoV-2^ binding hISG15 and Ub_2_ but also sites that may discriminate binding between these two substrates. Guided by these predictions we tested four mutants, two in the S2 site (F69A and E70A) and two in the S1 site (R166A and E167A) that have different predicted effects on binding to Ub_2_ and hISG15 (Fig. 5b, c). We again used MST to measure binding affinities for these alanine mutants to test their effect on PLpro^CoV-2^ binding to hISG15 and Ub_2_. Similarly to described in the PLpro^CoV-2^-C111S binding data (Supplementary Fig. 2a and Supplementary Table 1), we evaluated both 1:1 and 1:2 binding models determining that the 1:1 fits were sufficient to explain the binding profiles (Supplementary Fig. 13a, b and Supplementary Table 1). Consistent with the computational predictions, PLpro^CoV-2^-E167A has a dramatically reduced affinity for hISG15, Ub_2_ and Ub_1_ (Fig. 5f-h, blue curve and Fig. 5i, blue bars). This interaction formed between E167 and R153 of hISG15 or R42 of Ub is important and stabilizing (Supplementary Fig. 13c) and is observed in the crystal structure. By contrast, our predictions suggested that R166A should have a more moderate effect on binding to substrate as R166 forms hydrogen bond with similar geometries to N151 or Q49 of hISG15 and Ub_2_, respectively (Supplementary Fig. 13c). In the MST experiment, R166A has a larger effect on hISG15 binding compared to Ub_2_ or Ub_1_ (Fig. 5f-h, green curve and Fig. 5i, green bars). Our calculations of interface energetics also indicated that PLpro^CoV-2^ F69A should have a larger effect on interactions with hISG15 compared to Ub_2_, including the orientation observed in the distal Ub from the partial model of our structure (Supplementary Fig. 13). Indeed, we find that PLpro^CoV-2^-F69A alters binding to hISG15 significantly and only has a modest effect on Ub_2_ or Ub_1_ binding (Fig. 5f-h, orange curve and Fig. 5i, orange bars). A closer look at the F69 interacting residues reveals that hISG15 uses M23 to pack against the phenylalanine while the shorter side chain of I44 in Ub_2_ is more distant. Finally, we evaluated PLpro^CoV-2^-E70A binding to hISG15, Ub_2_ and Ub_1_. Our predictions suggested that E70A should only weakly decrease stability of PLpro^CoV-2^:hISG15 and PLpro^CoV-2^:Ub_2_ complexes (Fig. 5b and Supplementary Fig. 12) but the MST measurements show that PLpro^CoV-2^ E70A cannot bind well to Ub_2_ or Ub_1_ while binding to hISG15 is less affected (Fig. 5f-h, yellow curve and Fig. 5i, yellow bars). While this observation was not predicted correctly by the *in silico* alanine scan, inspection of the structures provides clues. In the PLpro^CoV-2^:hISG15 structure E70 is close to S22 but does not form any clear stabilizing interactions (Supplementary Fig. 13c); by contrast, E70 forms a long salt bridge to H68 and K6 in the PLpro^CoV-2^:Ub_2_ structure that may have been missed by the *in silico* analysis due to the larger distance (Supplementary Fig. 13c, > 3.9Å and 5Å, respectively). Coincidentally, K6 yielded heterogeneous cross-links to acidic residues, including E70, on the thumb domain of PLpro (Fig. 4d) suggesting that charge complementary interactions between the distal Ub and PLpro contribute to the tight binding between PLpro^CoV-2^ and Ub_2_.

Overall, our combined approach using *in silico* analysis of the interfaces and binding measurements of mutants uncovered exciting new means to discriminate binding between PLpro and hISG15 or Ub_2_, allowing future experiments in more physiological contexts to begin decoupling hISG15 and Ub-dependent effects on virulence and disease. Furthermore, our data suggest that distal domain binding of hISG15 and Ub_2_ to PLpro are determined by different types of interactions (i.e. nonpolar vs charge complementary electrostatics) thus the association or dissociation rates dictate binding that ultimately manifests as changes in dynamics of the distal domain of Ub (in Ub_2_) vs UBL (in hISG15) (Figure 6).

**Figure 6.**
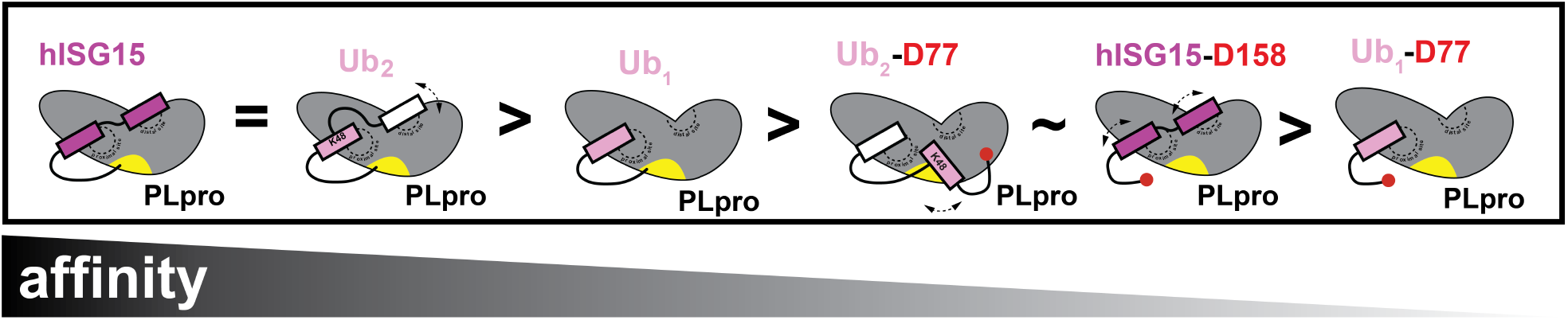
Dual domain-based model for PLpro recognition of K48-linked Ub_2_ and ISG15. Schematic representation of differences in binding of PLpro^CoV-2^ with hISG15 and ubiquitin species.

## DISCUSSION

The literature highlights that PLpro^CoV-1^ prefers binding to K48-linked polyubiquitin over ISG15^28^. Our work incited by recent studies^26,27^ reveals that PLpro^CoV-2^ recognizes human ISG15 and K48-Ub_2_ with very similar affinities, both in the nanomolar range. Sequence analysis suggests that the PLpro from these two viruses only vary at 8 amino acid positions at the substrate binding interface implicating only minor sequence changes responsible for improving binding of hISG15. Interestingly, both K48-Ub_2_ and ISG15 utilize two Ub/UBL domains to recognize and bind PLpro^CoV-2^ but our data show they do it differently. In ISG15 the two UBL domains are connected through a relatively short, likely more rigid peptide linker (DKCDEP in hISG15), and the C-terminal (proximal) UBL binds to S1 site while the N-terminal (distal) UBL binds to S2 site on PLpro^CoV-2^. The amino acid sequences of the proximal and distal UBLs are somewhat different and show distinct contacts that are required for tight binding. In K48-Ub_2_ two identical Ub units are connected through a flexible linker (RLRGG-K48)^33,34^ and contribute differently to binding. In the high affinity complex with PLpro the proximal Ub binds to the S1 site and the distal Ub binds to S2 site. The proximal Ub is very well ordered and shows multiple interactions with PLpro. The distal Ub interacts with PLpro differently. In the crystal structure this Ub domain is less ordered, suggesting multiple possible states. NMR data clearly show interactions of the distal Ub with PLpro, but presence of less occupied states cannot be excluded. K48-Ub_2_ also binds to PLpro with an altered register where the distal Ub binds to the S1 site while the proximal Ub occupies the S1’ site. This also supports how PLpro disassembles Ub_3_ by cleaving off the unit 3 and the fact that the binding curves of Ub_2_ and Ub_1_ to PLpro^CoV-2^ are explained better by presence of two binding events. Both hISG15 and the K48-Ub_2_ should be sensitive to mutations of S1 and S2 sites in PLpro, but because of different interaction modes, binding of the hISG15 and the K48-Ub_2_ substrates is likely to have different sensitivity to such mutations. Furthermore, evolutionary analysis of SARS-CoV-2 variants (Supplementary Fig. 14) highlights sequence variation at key sites including in the thumb domain, where we have shown that mutations differentially impact ISG15 and Ub_2_ specificity. Our engineered mutations uncovered nonpolar-vs electrostatics-driven distal UBL (F69) and Ub (E70) contacts in the PLpro^CoV-2^:hISG15 and PLpro^CoV-2^:Ub_2_ complexes, respectively, that likely underlie the differences in dynamics of the two substrates binding to the protease. These data also suggest that evolutionary variation in the PLpro sequence may alter substrate binding, potentially differentially dysregulating Ub and ISG15 processing but currently it is unknown how these alter disease outcomes. Our mutational data suggests that we may be able to engineer mutations that can shift the preference in substrate processing and directly test the contribution of each proteolytic activity in viral pathogenesis.

The structure of PLpro^CoV-1^:K48-Ub_2_ complex^28^ revealed both Ubs bound to the protease S1 and S2 binding sites, with proximal and distal Ubs connected via a non-hydrolyzable triazole linker (in lieu of the native isopeptide linkage) and the C-terminal tail covalently attached to the protease. This may rigidify the K48-Ub_2_ concealing the true interactions. There is no structure of the ISG15 bound to PLpro^CoV-1^ that could reveal their interactions. Our structure of PLpro^CoV-2^:hISG15 as well as the previous structure of PLpro^CoV-2^:mISG15 reveal a dual UBL domain recognition binding mode despite surprising species sequence variation at the UBL binding surfaces (Supplementary Fig. 1a-c). Our *in silico* alanine scan of the interfaces uncovered residues that may play central roles in stabilizing the ISG15 binding mode for PLpro^CoV-2^ and implicated V66 and F69 as being important for stabilizing the distal UBL domain of ISG15.

Much of the focus on understanding how sequence variation impacts pathogenicity, infectivity and virulence of SARS-CoV-2 has been centered on sequence changes in surface proteins such as the receptor-binding domain (RBD) of the spike protein which are essential for recognition of ACE2, virus entry into host cell and thus infectivity. Furthermore, there is concern that mutations in the viral receptors may overcome vaccines which were designed against an engineered prefusion “stabilized” conformation of RBD of the spike protein, particularly worrisome with emerging new variants like the Omicron BA.2 and Ontario WTD clade^41^. Therefore, additional SARS-CoV-2 life cycle steps must be explored, and appropriate key drug targets identified to expand treatment options.

Viral interference with host innate immune response is one of these steps of which ISG15 is integral. Several coronavirus Nsps have been shown to contribute to diminishing this complex response mechanism. Modeling of the protein interfaces suggests that the sequence variation between PLpro from SARS-CoV-1 and SARS-CoV-2 plays a role in recognition specificity of host factors. Furthermore, we also show that sequence variation within PLpro from 2.3 million SARS-CoV-2 isolates is overall distributed with some hotspots that mimic sequence variation observed between SARS-CoV-1 and SARS-CoV-2 (Supplementary Fig. 14). While we do not understand how differential recognition of Ub compared to ISG15 impacts pathogenicity and virulence of SARS coronaviruses, balance between dysregulation of the protective interferon response and ubiquitin-proteasome systems likely influences virus interference with the host defense mechanisms. Future work must be focused on understanding how protease specificity impacts pathogenicity. Furthermore, it remains unknown whether PLpro encodes additional specificity for the substrates that are linked to Ub or ISG15 modifications.

PLpro^CoV-2^ must recognize and process multiple substrates: polyproteins 1a and 1ab, polyUb and ISG15. It is also known to cleave several other human host proteins. All these substrates have common sequence recognition motif “LRGG”, however they differ in several ways. In coronavirus polyproteins and several host proteins PLpro cleaves a regular peptide bond. In K48-polyUb and ISG15-modified protein substrates the cleaved isopeptide bond is between the C-terminal carboxylate of Ub/UBL and a lysine side chain of Ub or other protein, but these latter substrates differ in how the Ubs or UBLs are linked. It is interesting that conformation of PLpro in complexes with Ub, hISG15 and mISG15 is very similar (0.7 Å rmsd). However, the substrates conformations differ.

What may be the biological implication of single vs dual domain recognition of Ub/UBL for hydrolysis of polyUb/ISG15 modifications? ISG15 is gene-coded fused dimer, it functions as a di-UBL and is attached covalently to proteins as such. Its specific removal by viral PLpro is also hard-wired to its dimer structure. Ubiquitin is different as it exists as a monomer and is added to a polyUb chain or other proteins in units of monomer. However, PLpro shows the highest affinity for Ub_2_ (or presumably longer chains) vs Ub_1_ and it most efficiently removes the proximal Ub (unit 3) from Ub_3_. Because PLpro binds single Ub less strongly, this suggests that in the cell proteins tagged with polyUb chains containing an odd number of Ubs may accumulate Ub_1_-substrate adducts, as shown in our cleavage studies.

PLpro^CoV-2^ binds both hISG15 and K48-Ub_2_ with high and similar affinity but shows weaker interactions with Ub_1_ or ISG15_prox/_ISG15_disal_. Our data also suggest the presence of lower affinity secondary binding events for hISG15 and K48-Ub_2_ which can be explained by alternate binding modes. However, the binding mode of these two substrates is quite different. Binding of hISG15 is defined by well-ordered proximal and distal UBLs bound to S1 and to S2 sites, respectively. This may be explained by a combination of a more rigid short (uncleavable) peptide linker between domains and the types of stabilizing interactions between the distal domain of Ub/UBL and PLpro.

In this mode hISG15 should be cleaved off substrate protein positioned at S1’ very efficiently and accumulate free ISG15 at high viral level. K48-Ub_2_ binds predominantly using proximal Ub domain to S1 site. The distal domain binds to S2 site but it is less ordered and assumes different states that still contribute to increased affinity. However, in order to be cleaved, the Ub_2_ substrate must switch to a different, lower affinity mode with the distal Ub bound to S1 site and the proximal Ub positioned at the S1’ site. Moreover, we noticed that the amino acid sequence of the substrates’ tail entering the active site (the “LXGG↓X” motif) also influenced rate of cleavage, with acidic residues (E/D) poorly tolerated next to the Gly residue, most likely disrupting ionization states of catalytic triad residues (Cys-His-Asp).

In summary, our findings pave the way to understand the interaction of PLpro with hISG15 and (poly)ubiquitin substrates and uncover binding heterogeneity that appears to decouple binding affinity from protease activity. Future experiments will focus on how sequence changes in PLpro can influence the distribution of primary and secondary binding sites of substrates. These experiments will be essential to decouple the different proteolytic activities (Nsps, polyUb and ISG15) and determine their contribution to viral pathogenesis.

## METHODS

### Gene cloning, protein expression and purification of WT and mutants of PLpro

The gene cloning, protein expression and purification were performed using protocols published previously^42^. Briefly, the Nsp3 DNA sequence corresponding to PLpro protease of SARS-CoV-2 was optimized for *E. coli* expression using the OptimumGene codon optimization algorithm followed by manual editing and then cloned directly into pMCSG53 vector (Twist Bioscience). The plasmids were transformed into the *E. coli* BL21(DE3)-Gold strain (Stratagene). *E. coli* cells harboring plasmids for SARS-CoV-2 PLpro WT and mutants (C111S; C111S,F69A; C111S,E70A; C111S,R166A; C111S,E167A and C111S,D286N) and ISG15 expression were cultured in LB medium supplemented with ampicillin (150 μg/ml).

For large-scale purification of WT and mutant PLpro^CoV-2^ constructs, 4 L cultures of LB Lennox medium were grown at 37 °C (200 rpm) in the presence of ampicillin 150 μg/ml. Once the cultures reached OD_600_ ∼1.0, the temperature setting was changed to 4 °C. When the bacterial suspensions cooled down to 18 °C they were supplemented with 0.5 mM IPTG and 40 mM K_2_HPO_4_ (final concentration). The temperature was set to 18 °C for 20 hours incubation. Bacterial cells were harvested by centrifugation at 7,000x g and cell pellets were resuspended in a 12.5 ml lysis buffer (500 mM NaCl, 5% (v/v) glycerol, 50 mM HEPES pH 8.0, 20 mM imidazole pH 8.0, 10 mM β-mercaptoethanol, 1 μM ZnCl_2_) per liter culture and sonicated at 120W for 5 minutes (4 sec ON, 20 sec OFF). The cellular debris was removed by centrifugation at 30,000x g for 90 minutes at 4 °C. The supernatant was mixed with 3 ml of Ni^2+^ Sepharose (GE Healthcare Life Sciences) which had been equilibrated with lysis buffer supplemented to 50 mM imidazole pH 8.0, and the suspension was applied on Flex-Column (420400-2510) connected to Vac-Man vacuum manifold (Promega).

Unbound proteins were washed out via controlled suction with 160 ml of lysis buffer (with 50 mM imidazole pH 8.0). Bound proteins were eluted with 15 ml of lysis buffer supplemented to 500 mM imidazole pH 8.0, followed by Tobacco Etch Virus (TEV) protease treatment at 1:25 protease:protein ratio. The solutions were left at 4 °C overnight. Size exclusion chromatography was performed on a Superdex 75 column equilibrated in lysis buffer. Fractions containing cut protein were collected and applied on a Flex-Column with 3 ml of Ni^2+^ Sepharose which had been equilibrated with lysis buffer. The flow through and a 7 ml lysis buffer rinse were collected. Lysis buffer was replaced using a 30 kDa MWCO filter (Amicon-Millipore) via 10X concentration/dilution repeated 3 times to crystallization buffer (20 mM HEPES pH 7.5, 150 mM NaCl, 1 μM ZnCl2, 10 mM DTT). The final concentration of WT PLpro^CoV-2^ was 25 mg/ml and C111S mutant was 30 mg/ml. For some NMR studies requiring longer measurement times we also utilized C111S,Y171H PLpro^CoV-2^ variant which showed similar binding properties to C111S variant but was more stable in the NMR buffer.

### Expression and purification of unlabeled and isotope labeled hISG15 and Ub

Human ISG15, as well as genes for the distal (hISG15_distal_) and proximal (hISG15_prox_) UBLs separately, were also synthesised and cloned directly into pMCSG53 vector. The UBL sequences included amino acids G2–L82 (hISG15_distal_) and L82–G157 (hISG15_prox_) of hISG15. These constructs were purified following the same protocol as for PLpro except that the buffers did not contain ZnCl2 and a 10 kDa MWCO filter was used for buffer exchange and concentration. The N-terminal polyhistidine tag was removed using TEV protease which left an additional serine/alanine at the N-terminus followed by an additional Ni-NTA step. The final concentration of hISG15 was 40 mg/ml. Unlabeled Ub variants were expressed in BL12(DE3) *E*.*coli* cells containing a helper pJY2 plasmid and purified as described elsewhere^29^. For expression of uniformly ^15^N-labeled Ubs, hISG15, hISG15_distal_, hISG15_prox_, as well as PLpro^CoV-2^ variants the cells were grown in minimal media containing ^15^NH_4_Cl as the sole source of nitrogen using methods previoulsy described in Varadan et al^29,31,43^.

### Synthesis of K48-polyUb chains

Ub_2_ and Ub_3_ chains were assembled from the respective recombinant Ub monomers using controlled chain synthesis catalyzed by Ub-activating E1 enzyme UBA1 and K48-specific Ub-conjugating E2 enzyme UBE2K (aka E2-25K) as detailed elsewhere^29,31,43^. Specifically, Ub variants bearing chain-terminating mutations, Ub-K48R and Ub-D77, were used to ensure that only Ub dimers are produced and to enable incorporation of ^15^N labeled Ub units at the desired distal (^15^N-Ub-K48R) or proximal (^15^N-Ub-D77) position in the resulting chain for NMR and MS studies. D77 was subsequently removed from the proximal Ub by Ub C-terminal hydrolase YUH1. Ub_3_ chains were made in a stepwise manner, by first removing D77 from the proximal Ub in Ub_2_ by YUH1 and subsequently conjugating this Ub_2_ to Ub-D77 using E1 and E2-25K to produce Ub_3_. The Ub_3_ chain for MS-based cleavage assays had Ub-K48R at the distal unit, ^15^N-labeled Ub at the endo unit, and Ub-D77 at the proximal unit, in order to allow unambiguous identification of the possible cleavage products by mass spectrometry. The correct masses of the synthesized Ub_2_ and Ub_3_ chains were confirmed using ESI-MS and SDS-PAGE.

### Microscale Thermophoresis binding assay

MST experiments were performed using NanoTemper Monolith NT.115 available in the Macromolecular Biophysics Resource core at UTSW and standard protocol was employed during analysis^44^. Ub_1_, Ub_1_-D77, K48-Ub_2_, K48-Ub_2_-D77, hISG15, hISG15-D158, hISG15_distal_ or hISG15_proximal_ were labeled with Cyanine5 NHS ester dye (Cy5) and titrated by a 1:1 serial dilution of PLpro^CoV-2^ (C111S; C111S,F69A; C111S,E70A; C111S,R166A and C111S,E167A mutants).

Obtained data were fit and analyzed in PALMIST 1.5.8 using 1:1 and 1:2 binding models and visualized in GUSSI 1.4.2^44,45^. All measurements were done in technical triplicates. To determine whether the 1:1 or 1:2 binding models were more suitable we calculated the probability of getting the *χ*^2^ improvement by chance using a t-test. The binding stoichiometry of the complexes was verified using cross-linking of complexes and visualized by SDS-PAGE. PALMIST and GUSSI software is freely available for academic users on UTSW MBR core’s website (https://www.utsouthwestern.edu/labs/mbr/software/). A summary <u>of MST fitting</u> parameters (K_d_s and errors) for 1:1 and 1:2 binding data are shown in Supplementary Table 1.

### PLpro cleavage assay

K48-Ub_3_, K48-Ub_2,_ K48-Ub_2_-D77 and ISG15-Nsp2, ISG15-Nsp3 and ISG15-Nsp4 cleavage reactions were performed at 20 °C in 20 mM Tris buffer (pH 7.52) containing 100 mM NaCl, 10 mM DTT, 1 μM ZnCl_2_. The initial volume was 350 μl and contained 20 μM of Ub_3_. Upon addition of 0.5 μM PLpro, equal amounts (10 μl) of reaction samples were aliquoted out at given time points, mixed with equal volume of SDS load buffer and immediately placed in a water bath at 70 °C for 5 minutes to stop the reaction. The samples were then loaded onto 15% urea polyacrylamide gel and resolved using SDS-PAGE. For mass spectrometry analyses cleavage reactions were performed on ice, the samples were buffer exchanged into autoclaved RO water and concentrated to 50 μl volume prior to analysis. The data shown here were obtained on Bruker Maxis-II ultra-high resolution Q-TOF mass spectrometer available at University of Maryland. Peaks were isolated during separation and analyzed using MagTran 1.03 b2.

### Protein crystallization

Crystallizations were performed with the protein-to-matrix ratio of 1:1 using the sitting drop vapor-diffusion method with the help of the Mosquito liquid dispenser (TTP LabTech) in 96-well CrystalQuick plates (Greiner Bio-One). MCSG1, MCSG2, MCSG3, and MCSG4 (Anatrace), Index (Hampton Research) and Wizard 1&2 (Jena Bioscience) screens were used at 16 °C. The PLpro^CoV-2^-C111S:hISG15 complex (13 mg/ml) crystallized in Index E11 (0.02 M MgCl2, 0.1 M HEPES pH 7.5, 22% (w/v) polyacrylic acid sodium salt 5,100). For the PLpro^CoV-2^-C111S:Ub2-K48R,D77 complex (11 mg/ml), crystals appeared in MCSG-2 F11 and were improved in hanging drops with a protein-to-matrix ratio of 2:1 in 0.2 M disodium tartrate, 15 % (w/v) PEG3350 after seeding with 1/10 volume of PLpro^CoV-2^-C111S:Ub2-K48R, D77 microcrystals. Crystals of the PLpro^CoV-2^C111S,D286N:Ub2-K48R complex were obtained in 0.2 M disodium tartrate, 15 % (w/v) PEG3350 (as above). The Ub_2_ protein crystallized in Pi-PEG D1 (50 mM acetate buffer pH 4.8, 8.6% PEG 2000 MME, 17.1% PEG 400). The hISG15 protein crystallized in MCSG1 G2 (40 mM potassium phoshate, 16% PEG 8000, 20% glycerol). Crystals selected for data collection were washed in the crystallization buffer supplemented with 25% glycerol and flash-cooled in liquid nitrogen.

### Data collection, structure determination, and refinement

Single-wavelength X-ray diffraction data were collected at 100 K temperature at the 19-ID beamline of the Structural Biology Center at the Advanced Photon Source at Argonne National Laboratory using the program SBCcollect. The diffraction images were recorded on the PILATUS3 × 6M detector at 12.662 keV energy (0.9792 Å wavelength) using 0.3° rotation and 0.3 sec exposure. The intensities were integrated and scaled with the HKL3000 suite^46^. Intensities were converted to structure factor amplitudes in the truncate program^47^ from the CCP4 package^48^. The structures were determined by molecular replacement using HKL3000 suite incorporating the program MOLREP^49–51^. The coordinates of PLpro^CoV-2^ in complex with ubiquitin propargylamide (PDB id: 6XAA) and PLpro^CoV-2^-C111S with mISG15 (PDB id: 6YVA) were used as the starting models for PLpro^CoV-2^-C111S:K48-Ub_2_ and PLpro^CoV-2^-C111S:hISG15 structure solutions, respectively. For Ub_2_ and hISG15 proteins, the structures of PLpro^CoV-1^ bound to a K48-Ub_2_ activity based probe (PDB id: 5E6J) and ISG15 (PDB id: 1Z2M) were used as the starting models. The initial solutions were refined, both rigid-body refinement and regular restrained refinement by REFMAC program^52^ as a part of HKL3000. Several rounds of manual adjustments of structure models using COOT^53^ and refinements with REFMAC program^52^ from CCP4 suite^48^ were done.The stereochemistry of the structure was validated with PHENIX suite^54^ incorporating MOLPROBITY^55^ tools. A summary of data collection and refinement statistics is given in Supplementary Table 2.

### NMR data collection and analysis

NMR measurements were performed at 25 °C on Avance III 600 MHz and 800 MHz Bruker NMR spectrometers equipped with cryoprobes. The data were processed using Topspin (Bruker) and analyzed using Sparky 3.190^56^. NMR signal assignments for hISG15 were obtained from Biological Magnetic Resonance Data Bank (BMRB Entry ID 5658) and adjusted to the temperature and buffer conditions used in our studies.

The protein samples for NMR measurements were prepared in 50 mM HEPES buffer or in 20 mM Tris buffer, both at pH 7.42 and containing 100 mM NaCl, 1 mM TCEP, 1 μM ZnCl_2_, 0.2% (w/v) NaN_3_, and 10% (v/v) D_2_O. Binding studies by NMR were carried out by adding pre-calculated amounts of unlabeled PLpro^CoV-2^-C111S to ^15^N-labeled hISG15 or K48-Ub_2_ (with either distal or proximal Ub ^15^N labeled) up to ∼1:1 molar ratio, or 2:1 molar ratio to Ub_1_ and monitoring changes in 2D ^1^H-^15^N SOFAST-HMQC and/or ^1^H-^15^N TROSY as well as 1D ^1^H spectra. The initial binding studies were performed in HEPES buffer and subsequently repeated in Tris buffer, both produced similar results. Reciprocal titrations were performed by adding unlabeled Ub_2_, Ub_1_, or hISG15 (or D158-extended) to ^15^N-labeled PLpro^CoV-2^-C111S or PLpro^CoV-2^-C111S,Y171H in a 1.5:1 (Ub_2_, hISG15:PLpro) or up to 8:1 (Ub_1_:PLpro) molar ratio_. 15_N-labeled PLpro^CoV-2^-C111S,Y171H was primarily used for lengthy TROSY experiments as this variant proved to be more stable than PLpro^CoV-2^-C111S,Y171H in the NMR buffer. Both PLpro^CoV-2^ variants had very similar NMR spectra and essentially identical signal perturbations upon binding of all the substrates tested (see Supplementary Fig. 8).

The protein concentrations in NMR studies in Tris buffer were as follows: 150:150 μM for hISG15:PLpro^CoV-2^, 180:270 μM for hISG15-D158:PLpro^CoV-2^, 83:104 μM for ^15^N-distal Ub_2_:PLpro, 200:200 μM for ^15^N-proximal Ub_2_:PLpro^CoV-2^, 152.8:305.9 μM for Ub_1_:PLpro^CoV-2^, 125:250 μM for Ub_1_-D77:PLpro^CoV-2^. Experiments with ^15^N-PLpro^CoV-2^ used 301.5:201.5 μM hISG15:PLpro, 225:150 μM hISG15-D158:PLpro, 137.5:110 μM Ub_2_:PLpro, 404:202 μM Ub_2_-D77:PLpro, up to 1140:141 μM Ub_1_:PLpro, 1000:125 μM Ub_1_-D77:PLpro. For NMR measurements in HEPES buffer, the concentrations were 115:115 μM for hISG15: PLpro^CoV-2^; 71:142 μM for ^15^N-distal Ub_2_: PLpro^CoV-2^; 75:150 μM for ^15^N-proximal Ub_2_: PLpro^CoV-2^; and 81:151 μM for Ub_1_:PLpro^CoV-2^.

### Cross-linking mass spectrometry analysis

Our group has developed standardized protocols for cross-linking and data analysis of samples. For complexes between PLpro^CoV-2^and Ub_1_, Ub_1_-D77, K48-Ub_2_, K48-Ub_2_-D77 or hISG15, we incubated the protease with the substrate at a 1:4 molar ratio for 1 hour at 25 °C. For disuccinimidyl suberate (DSS)^57^ reactions, samples were cross-linked with a final 1 mM DSS (DSS-d_0_ and -d_12_, Creative Molecules) for three minutes at 37 °C while shaking at 350 rpm. For sulfonyl fluoride (SuFEx)^58^ reactions, samples were cross-linked with a final 0.658 mM SuFEx (a kind gift from Dr. William DeGrado, UCSF) for one hour at 37 °C while shaking at 350 rpm. For DMTMM^38^ reactions, samples were cross-linked with a final 43 mM DMTMM (Sigma-Aldrich) for 15 minutes at 37 °C while shaking at 600 rpm. All cross-linking reactions were quenched with 172 mM (4 times excess) ammonium bicarbonate for 30 minutes at 37 °C while shaking at 350 rpm. Samples were resolved on SDS-PAGE gels (NuPAGE™, 4 to 12%, Bis-Tris, 1.5 mm) and, for DMTMM cross-linker set of samples, bands corresponding to PLpro^CoV-2^:substrate heterodimers were gel-extracted following standard protocols^59^. For glutaraldehyde cross-linking, we used 1:4, 1:1 and 1:0.25 ratios of PLpro:substrate using 6 μM protease in all reactions. Samples were preincubated at 25 °C for 15 minutes followed by addition of 0.05% glutaraldehyde (Sigma) for 1 minute and quenched with Tris pH 8 to a final concentration of 0.2 M. Cross-linked species and control reactions were resolved by SDS-PAGE. Cross-linked samples for XL-MS were digested by 1:50 (m/m) trypsin (Promega) overnight at 37 °C while shaking at 600 rpm. 2% (v/v) formic acid was added to acidify the reaction and further purified by reversed-phase Sep-Pak tC18 cartridges (Waters), next flash frozen in liquid nitrogen and lyophilized. The dried samples were resuspended in water/acetonitrile/formic acid (95:5:0.1, v/v/v) to a final concentration of approximately 0.5 μg/μl. 2 μl of each was injected into Eksigent 1D-NanoLC-Ultra HPLC system coupled to a Thermo Orbitrap Fusion Tribrid system at the UTSW Proteomics core.

The mass spectrometry data were analyzed by in-house version of xQuest 2.1.5 pipeline^60^. Thermo RAW data files were first converted to open .mzXML format using msconvert (proteowizard.sourceforge.net). The mass spectra across replicates yielded similar intensities. Search parameters were set based on DMTMM as the cross-link reagent as follows: maximum number of missed cleavages = 2, peptide length = 5-50 residues, fixed modifications = carbamidomethyl-Cys (mass shift = 57.02146 Da), variable modification = oxidation of methionine (mass shift = 15.99491 Da), mass shift of cross-linker = -18.010595 Da, no monolink mass specified, MS1 tolerance = 15 ppm, and MS2 tolerance = 0.2 Da for common ions and 0.3 Da for cross-link ions; search in enumeration mode. Next, in-house shell script was employed to identify cross-links between lysines and acidic residues. FDRs were estimated by xProphet^61^ to be 9.8% - 77.8%. For each experiment, five replicate data sets were compared and only cross-link pairs that appeared in all data sets (PLpro^CoV-2^:K48-Ub_2_ and PLpro^CoV-2^:Ub_1_) or at least in four data sets (PLpro^CoV-2^:hISG15, PLpro^CoV-2^:K48-Ub_2_-D77) were used to generate a consensus data set (Source Data 1).

### Modeling of alternate Ub binding sites on PLpro^CoV-2^

To build an ensemble of alternate Ub binding sites on PLpro^CoV-2^ outside of the canonical proximal domain on the S1 site, we employed docking procedure that combined a geometric restraint between K48 of the immobile proximal Ub to the C-terminus of the mobile Ub to sample alternate S2 binding sites. In the alternative scenario, we employed a geometric restraint between C-terminus of the immobile proximal Ub to K48 of the mobile Ub to sample alternate S1’ binding sites below the active sites. An initial conformation of the PLpro^CoV-2^ bound to Ub in the proximal site was built from our structure (PDB id: 7RBR) and converted into a single chain. We next added a mobile Ub as a second chain and produced over 5000 low-resolution centroid mode models employing two different geometric restraints that sampled alternate S2 and S1’ binding sites. Each structure was minimized and the total energy of the PLpro^CoV-2^:Ub_2_ complexes was plotted as a function of a sum distances of Cβ-Cβ from the experimental cross-links using an in-house script.All simulations were performed with RosettaDock protocol^62^ as a part of Rosetta 3.13 suite and ran on UTSW’s BioHPC computing cluster. All plots were generated with GraphPad Prism 9. Images were created using PyMOL 2.5.1.

### Energetic analysis of PLpro^CoV-2^ in complex with hISG15 and K48-Ub_2_

Models of PLpro^CoV-2^ bound to two different substrates, hISG15 and K48-Ub_2_, that were used in the subsequent *in silico* alanine scan were prepared as follows. For the PLpro^CoV-2^:hISG15 complex, we used a heterodimer conformation derived from our crystal structure (PDB id: 7RBS). To create the complex between PLpro^CoV-2^ and K48-Ub_2_, we used the conformation and binding mode of K48-Ub_2_ bound to PLpro^CoV-1^ (PDB id: 5E6J) as a template^28^. Briefly, our PLpro^CoV-2^ (PDB id: 7UV5) was aligned to PLpro^CoV-1^ bound to K48-Ub_2_ to produce a tentative model of PLpro^CoV-2^:K48-Ub_2._ As a control, a model of PLpro^CoV-2^ with single proximal Ub visible in our density was also analyzed. Next, we applied a relax protocol in Rosetta for both complexes: PLpro^CoV-2^:K48-Ub_2_ and PLpro^CoV-2^:hISG15. To guarantee that each instance of relax is being run with different randomizations, groups of nstruct were run with different, randomly generated seeds using random.org. From 100 total structures (4×25 nstruct for computational efficiency) for each heterocomplex, the lowest energy structure was identified and used in further steps. The list of PLpro residues that may be engaged in interacting with its substrate was created by identifying PLpro residues within 4.0 Å of either hISG15 or K48-Ub_2_. The union list of interacting residues identified with heterocomplexes PLpro^CoV-2^:hISG15 and PLpro^CoV-2^: K48-Ub_2_ was used in the next step. For PLpro^CoV-2^ in complex with hISG15 or K48-Ub_2_ 51 positions were used to describe the combined interface. Selection of common interface residues was carried out in PyMOL 2.5.1. Flex ΔΔG protocol was used as described previously^39^. The code is available on GitHub: https://github.com/Kortemme-Lab. Briefly, selected interacting residues were mutated to alanines. Parameters were used (all default settings): nstruct = 35, max_minimization_iter = 5000, abs_score_convergence thresh = 1.0, number_backrub_trials = 35000, and to enable earlier time points backrub_trajectory_stride was set to 7000. The ΔΔG_binding_ score for the last iteration is shown in the Results. These simulations were performed using BioHPC computing cluster at UT Southwestern Medical Center. The results, in raw REU (Rosetta energy units), are shown as a heat-map with ΔΔG_binding_ values but also as the difference between the ΔΔG_binding_ from hISG15 compared to K48-Ub_2_. The plots were made using GraphPad Prism 9 and mapped onto the protease structure using PyMOL 2.5.1. The relax protocol and Flex ddG used Rosetta v3.13 and v3.12, respectively.

### Sequence comparison of Ub, ISG15 and sequence variation across PLpro^CoV-2^ in SARS-CoV-2

Alignments were produced in Clustalo^63^ and visualized in Seaview^64^. Sequence identity between Ub_2_, hISG15 and mISG15 was calculated using Blast^65^. PLpro^CoV-2^ sequence variation from 2.3 million sequences (as of October 18, 2021) was derived from the coronavirus3D database^66^. The per residue mutational frequencies were mapped onto a PLpro^CoV-2^ structure in the context of a bound K48-Ub_2_ or hISG15.

### Reporting summary

Further information on research design is available in the <u>Nature Research Reporting</u> Summary linked to this paper.

### Statistics and Reproducibility

The results included in this manuscript can be reproduced by following protocols and using materials described in Methods.

### Data availability

The structural datasets generated during the current study are available in the Protein Data Bank repository (https://www.rcsb.org/) under PDB id: 7RBR for PLpro^CoV-2^, C111S mutant, in complex with K48-Ub_2_; PDB id: 7RBS for PLpro^CoV-2^, C111S mutant, in complex with human ISG15; PDB id: 7UV5 for PLpro^CoV-2^, C111S,D286N mutant, in complex with K48-Ub_2_; PDB id: 7S6P for human ISG15 alone, and PDB id: 7S6O for K48-Ub_2_ alone. Diffraction images are available on server in Dr. W. Minor laboratory https://proteindiffraction.org. Plasmid for expression PLpro^CoV-2^ C111S mutant (NR-52897 Vector pMCSG53 Containing the SARS-Related Coronavirus 2, Wuhan-Hu-1 Papain-Like Protease) is available in the NIH the BEI Resources Repository (https://www.niaid.nih.gov/research/bei-resources-repository). All MST, raw cross-linking mass spectrometry and ΔΔG_calc_ data are available in Source Data 1. Raw MS data used for the XL-MS analysis has been deposited in the MassIVE database under the accession number MSV000091075.

## CODE AVAILABILITY

All ΔΔG_calc_ calculations were performed using the flex_ddg protocol (available https://github.com/Kortemme-Lab/flex_ddG_tutorial) in Rosetta v3.12 (https://www.rosettacommons.org).

## ACKNOWLEDGEMENTS

We thank the members of the SBC at Argonne National Laboratory, especially Darren Sherrell, Krzysztof Lazarski and Alex Lavens for their help with setting beamline and data collection at beamline 19-ID. We thank Chad Brautigam in the UTSW Macromolecular Biophysics Resource core for help with MST data analysis. We also appreciate the help of the Proteomics Core Facility at UTSW. Funding for this project was provided in part by federal funds from the National Institute of Allergy and Infectious Diseases, National Institutes of Health, Department of Health and Human Services, under Contract HHSN272201700060C and 75N93022C00035 (AJ). The use of SBC beamlines at the Advanced Photon Source is supported by the U.S. Department of Energy (DOE) Office of Science and operated for the DOE Office of Science by Argonne National Laboratory under Contract No. DE-AC02-06CH11357. This research was supported by NIH grant GM065334 to D.F. NMR experiments were performed at UMD on instruments supported in part by NSF grant DBI1040158, and the instrument for mass spectrometry measurements was supported by NSF grant CHE-2018860. We acknowledge Dr. Yue Li at UMD for help with Maxis-II Q-TOF measurements. EE was supported by Undergraduate Maryland Summer Scholarship. LAJ is supported by an Effie Marie Cain Scholarship in Medical Research. PMW is supported by an O’Donnell Brain Institute Pilot grant.

## AUTHOR CONTRIBUTIONS

AJ, LAJ and DF initiated the project. RJ cloned, expressed and purified the first batch of protein, ME continued with different constructs, mutations and complexes. CT purified and crystallized protein and complexes with ligands and with JO collected diffraction data. JO determined, refined and together with KM analyzed structures. BTL and EE synthesized all Ub chains, expressed and purified all proteins for NMR studies and carried out NMR studies. BTL and CT performed Ub_3_ cleavage reactions and their MS analysis. PMW carried out all the MST binding, XL-MS and docking experiments and analysis. ΔΔG calculations and analysis were carried out by PMW and VM. Finally, DF, AJ, and LJ conceived of and directed the research as well as wrote the manuscript.

