## Supplementary Information File for "Dual domain recognition determines SARS-CoV-2 PLpro selectivity for human ISG15 and K48-linked di-ubiquitin"

**Source Data 1. Summary of raw cross-linking mass spectrometry, MST, and $\Delta\Delta G_{calc}$ data for $PLpro^{CoV\text{-}2}$:substrate complexes.**

**Supplementary Table 1.** Estimated binding constants and their associated confidence intervals from MST binding data fit to 1:1 and 2:1 binding models.

| | | 1:1 fitting model[a] | | 2:1 fitting model[a] | | | |
|---|---|---|---|---|---|---|---|
| **PLpro construct** | **Substrate** | $K_D$ | **68.3% ESP (error-surface projection) confidence interval** | $K_{D1}$ | **68.3% ESP (error-surface projection) confidence interval** | $K_{D2}$ | **68.3% ESP (error-surface projection) confidence interval** |
| C111S | hISG15 | 0.5[b] | [0.2, 1.5] | 0.09 | [0.03, 0.18] | 40 | [10, 350] |
| | $Ub_2$ (G76) | 0.04 | [0.01, 0.09] | 0.07 | [0.04, 0.11] | 170 | [20, N.D.[c]] |
| | $Ub_1$ (G76) | 24 | [12, 49] | 0.17 | [0.05, 0.48] | 70 | [50, 130] |
| | $Ub_2$ (D77) | 39 | [23, 73] | 0.6 | [0.1, 2.2] | 120 | [70, 470] |
| | $Ub_1$ (D77) | 130 | [90, 200] | N.D. | N.D. | N.D. | N.D. |
| | hISG15 (D158) | 6.1 | [4.6, 8.2] | 1.9 | [0, 4.9] | N.D. | N.D. |
| | $hISG15_{distal}$ | 90 | [50, 150] | N.D. | N.D. | N.D. | N.D. |
| | $hISG15_{proximal}$ | 90 | [30, N.D.] | N.D. | N.D. | N.D. | N.D. |
| C111S,F69A | hISG15 | 13 | [6, 29] | 0.3 | [N.D., 1.4] | 70 | [30, N.D.] |
| | $Ub_2$ (G76) | 1.6 | [1.2, 2.1] | 0.1 | [N.D., 0.7] | 3.5 | [1.9, 11.4] |
| | $Ub_1$ (G76) | 54 | [41, 72] | 8 | [1, 24] | 290 | [90, N.D.] |
| C111S,E70A | hISG15 | 2.9 | [1.1, 7.7] | 0.17 | [0.06, 0.42] | 34 | [16, 108] |
| | $Ub_2$ (G76) | 6 | [4, 10] | 0.5 | [0.1, 1.6] | 18 | [10, 42] |
| | $Ub_1$ (G76) | 43 | [30, 62] | 20 | [N.D., 140] | N.D. | N.D. |
| C111S,R166A | hISG15 | 15 | [8, 29] | 0.4 | [0.1, 0.9] | 50 | [30, 110] |
| | $Ub_2$ (G76) | 2.7 | [2, 3.5] | N.D. | N.D. | 3 | [1.9, 8.3] |
| | $Ub_1$ (G76) | 3.9 | [3.2, 4.6] | 1 | [0.2, 3.2] | 9 | [5, N.D.] |
| C111S,E167A | hISG15 | 42 | [29, 64] | 5 | [2, 9] | 600 | [100, N.D.] |
| | $Ub_2$ (G76) | 28 | [19, 44] | 0.14 | [0.03, 0.91] | 47 | [31, 80] |
| | $Ub_1$ (G76) | 70 | [51, 102] | 4 | [1, N.D.] | 140 | [60, N.D.] |

[a] Fitting of data to 1:1 and 1:2 binding model in PALMIST[1,2].

[b] All values in μM.

[c] Not determined.

**Supplementary Table 2.** Data processing and refinement statistics.

| Structure | **PLpro $^{CoV-2}$-C111S:K48-Ub$_2$** | **PLpro$^{CoV-2}$-C111S,D286N: K48-Ub$_2$** | **PLpro $^{CoV-2}$-C111S:hISG15** | **K48-Ub$_2$** | **hISG15** |
|---|---|---|---|---|---|
| **Data processing** | | | | | |
| Space group | C2 | C2 | C222$_1$ | P2$_1$ | P2$_1$ |
| Cell dimensions | | | | | |
| *a, b, c* (Å) | 148, 50, 72 | 148, 50, 72 | 154, 221. 233 | 24. 57, 46 | 52, 135, 78 |
| *α, β, γ* (°) | 90, 112, 90 | 90, 112, 90 | 90, 90, 90 | 90, 93, 90 | 90, 108, 90 |
| Resolution range (Å)[a] | 47.1-1.88 (1.91–1.88) | 36.7-1.45 (1.48–1.45) | 49.0-2.98 (3.08-2.98) | 46.1-1.25 (1.27–1.25) | 46.5-2.15 (2.19-2.15) |
| Unique reflections[a] | 38,678 (1,498) | 85,733 (4248) | 78,254 (3,893) | 28,558 (576) | 54,460 (2,545) |
| R-merge[b] | 0.106 (0.929) | 0.075 (1.24) | 0.214 (2.18) | 0.072 (0.50) | 0.070 (0.946) |
| Mean I/sigma(I) | 13.3 (1.10) | 25.3 (1.14) | 12.6 (1.23) | 21.2 (1.79) | 21.2 (1.12) |
| CC1/2[c] | 0.986 (0.520) | 0.996 (0.440) | 0.985 (0.364) | 0.994 (0.624) | 0.991 (0.511) |
| Completeness (%) | 96.1 (74.2) | 99.4 (99.2) | 99.9 (100) | 84.7 (34.9) | 98.3 (91.6) |
| Redundancy | 3.2 (2.2) | 5.4 (4.5) | 6.9 (6.7) | 3.2 (2.1) | 3.4 (2.8) |
| Wilson B-factor (Å$^2$) | 31.8 | 22.0 | 54.3 | 11.3 | 43.5 |
| **Refinement** | | | | | |
| Resolution range (Å) | 47.1-1.88 | 36.7-1.45 | 49.0-2.98 | 46.1-1.25 | 46.5-2.15 |
| Reflections work/test | 36,778 (1,900) | 81,507 / 4,226 | 74,428 (3,786) | 27,176 (1,363) | 51,806 (2,625) |
| $R_{work}/R_{free}$ | 0.187 / 0.228 | 0.136 / 0.179 | 0.195 / 0.236 | 0.129 / 0.177 | 0.221 /0.268 |
| RMSD (bonds) (Å) | 0.010 | 0.010 | 0.004 | 0.009 | 0.010 |
| RMSD (angles) (°) | 1.59 | 1.52 | 1.28 | 1.58 | 1.61 |
| Number of TLS groups | 2 | - [e] | 10 | - [e] | 6 |
| Total protein residues | 471 | 470 | 2,375 | 153 | 942 |
| **Number of atoms** | | | | | |
| Protein | 3,122 | 3,497 | 18,423 | 1,286 | 7,000 |
| Ligand / ion | 11 | 17 | 5 | 8 | - |
| Water | 169 | 355 | 54 | 141 | 119 |
| **B-factors** | | | | | |
| Average B-factor (Å$^2$) | 46.8 | 40.2 | 81.9 | 19.2 | 58.0 |

| | | | | | |
|---|---|---|---|---|---|
| Protein | 46.8 | 39.5 | 81.9 | 17.6 | 58.3 |
| Ligand / ion | 53.8 | 37.1 | 142.7 | 31.8 | - |
| Water | 46.8 | 46.9 | 52.8 | 33.3 | 44.8 |
| **MolProbity validation** | | | | | |
| Rotamer outliers (%)[d] | 0.87 | 0.76 | 1.62 | 0.65 | 5.67 |
| Clashscore[d] | 3.68 | 2.68 | 1.83 | 2.21 | 2.47 |
| Ramachandran outliers (%)[d] | 0.00 | 0.00 | 0.04 | 0.00 | 0.00 |
| MolProbity score[d] | 1.40 | 1.06 | 1.48 | 1.00 | 1.60 |
| **PDB ID** | 7RBR | 7UV5 | 7RBS | 7S6O | 7S6P |

[a] Values in parentheses correspond to the highest resolution shell.

[b] R-merge = $\Sigma_h \Sigma_j |I_{hj} - \langle I_h \rangle| / \Sigma_h \Sigma_j I_{hj}$, where $I_{hj}$ is the intensity of observation j of reflection h.

[c] As defined by Karplus and Diederichs[3].

[d] As defined by Molprobity[4].

[e] Anisotropic refinement of temperature factors (no TLS groups).

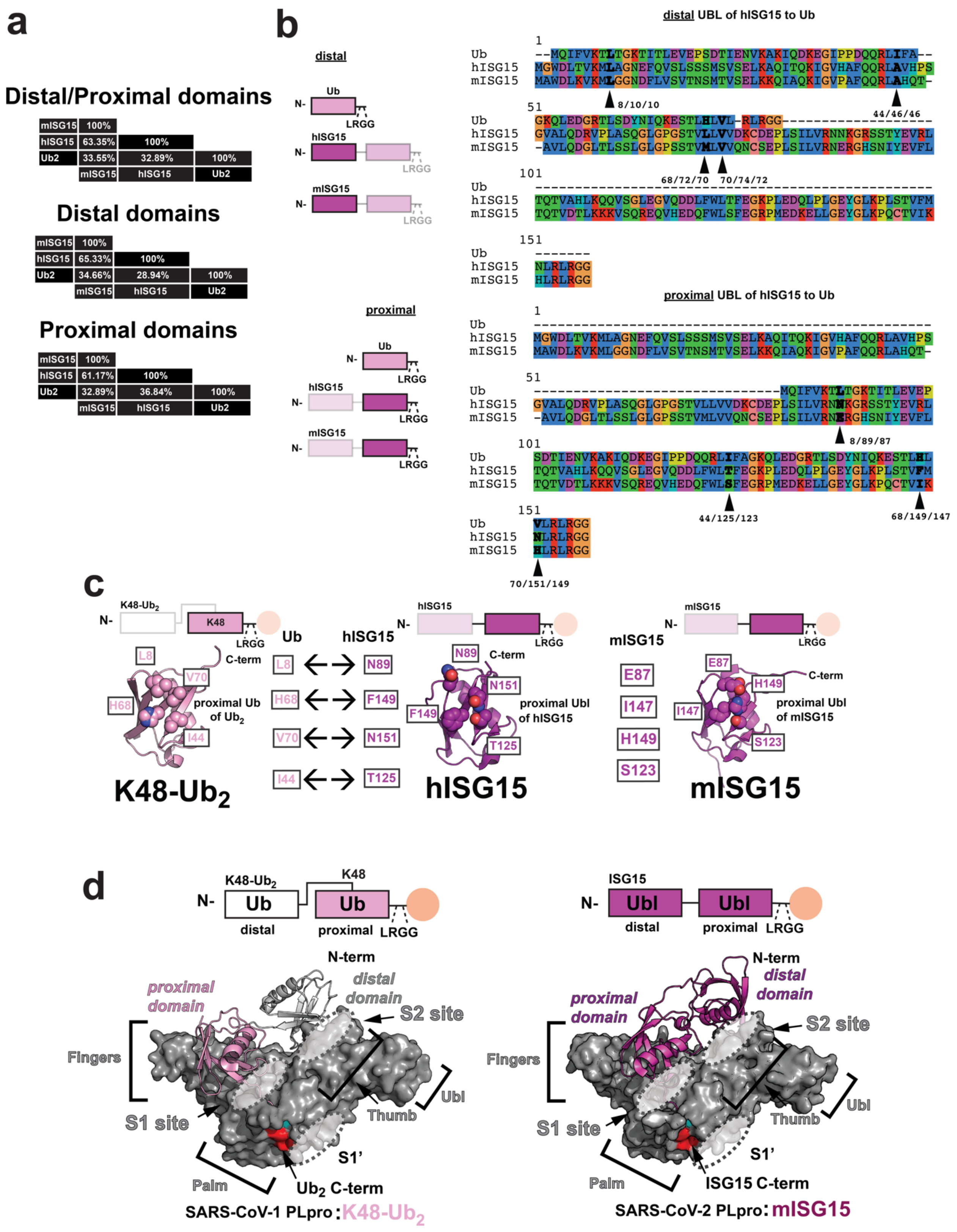
a
Distal/Proximal domains
mISG15 100%
hISG15 63.35% 100%
Ub2 33.55% 32.89% 100%
mISG15 hISG15 Ub2
Distal domains
mISG15 100%
hISG15 65.33% 100%
Ub2 34.66% 28.94% 100%
mISG15 hISG15 Ub2
Proximal domains
mISG15 100%
hISG15 61.17% 100%
Ub2 32.89% 36.84% 100%
mISG15 hISG15 Ub2
b
distal
Ub
hISG15
mISG15
LRGG
proximal
distal UBL of hISG15 to Ub
Ub --MQIFVKTLTGKTITLEVEPSDTIENVKAKIQDKEGIPPDQQRLIFA--
hISG15 MGWDLTVKMLAGNEFQVSLSSSMSVSELKAQITQKIGVHAFQQRLAVHPS
mISG15 MAWDLKVKMLGGNDFLVSVTNSMTVSELKKQIAQKIGVPAFQQRLAHQT-
8/10/10
44/46/46
Ub GKQLEDGRTLSDYNIQKESTLHLVL-RLRGG
hISG15 GVALQDRVPLASQGLGPGSTVLLVVDKCDEPLSILVRNNKGRSSTYEVRL
mISG15 -AVLQDGLTLSSLGLGPSSTVMLVVQNCSEPLSILVRNERGHSNIYEVFL
68/72/70
70/74/72
hISG15 TQTVAHLKQQVSGLEGVQDDLFWLTFEGKPLEDQLPLGEYGLKPLSTVFM
mISG15 TQTVDTLKKKVSQREQVHEDQFWLSFEGRPMEDKELLGEYGLKPQCTVIK
hISG15 NLRLRGG
mISG15 HLRLRGG
proximal UBL of hISG15 to Ub
hISG15 MGWDLTVKMLAGNEFQVSLSSSMSVSELKAQITQKIGVHAFQQRLAVHPS
mISG15 MAWDLKVKMLGGNDFLVSVTNSMTVSELKKQIAQKIGVPAFQQRLAHQT-
Ub MQIFVKTLTGKTITLEVEP
hISG15 GVALQDRVPLASQGLGPGSTVLLVVDKCDEPLSILVRNMKGRSSTYEVRL
mISG15 -AVLQDGLTLSSLGLGPSSTVMLVVQNCSEPLSILVRNERGHSNIYEVFL
8/89/87
Ub SDTIENVKAKIQDKEGIPPDQQRLIFAGKQLEDGRTLSDYNIQKESTLH
hISG15 TQTVAHLKQQVSGLEGVQDDLFWLTFEGKPLEDQLPLGEYGLKPLSTVFM
mISG15 TQTVDTLKKKVSQREQVHEDQFWLSFEGRPMEDKELLGEYGLKPQCTVIK
44/125/123
68/149/147
Ub VLRLRGG
hISG15 NLRLRGG
mISG15 HLRLRGG
70/151/149
c
K48-Ub2
K48
hISG15
mISG15
L8
V70
H68
I44
C-term
proximal Ub of Ub2
Ub
hISG15
L8 N89
H68 F149
V70 N151
I44 T125
N89
N151
F149
T125
proximal Ubl of hISG15
E87
I147
H149
S123
proximal Ubl of mISG15
K48-Ub2
hISG15
mISG15
d
K48-Ub2
Ub
distal
proximal
LRGG
ISG15
Ubl
N-term
proximal domain
distal domain
S2 site
Fingers
Thumb
Ubl
S1 site
S1'
Palm
Ub2 C-term
ISG15 C-term
SARS-CoV-1 PLpro : K48-Ub2
SARS-CoV-2 PLpro : mISG15

**Supplementary Figure 1. PLpro architecture and substrate recognition.** (**a**) Sequence identities between $Ub_2$, mISG15 and hISG15 for full sequences and individual domains: distal (N-terminal) and proximal (C-terminal) alone. (**b**) Sequence alignments of distal and proximal Ubs and UBLs of hISG15 and mISG15. Ub and ISG15 are shown in cartoon representation and are colored pink and magenta. Ub hydrophobic patch residues (L8, I44, H68, V70) important for Ub recognition and the corresponding residue positions in the proximal and distal UBL domains of hISG15 and mISG15 are highlighted by arrows. (**c**) Key residues that define the hydrophobic/nonpolar patch on Ub and ISG15 (proximal UBL domain) are shown in spacefill representation and highlighted with arrows. (**d**) Binding mode of K48-$Ub_2$ to $PLpro^{CoV\text{-}1}$ (left) and mISG15 to $PLpro^{CoV\text{-}2}$ (right). PLpro is shown in surface representation and colored gray. K48-$Ub_2$ is shown in cartoon representation and colored white and pink for the proximal and distal Ubs, respectively. Following the established terminology[5,6] for $Ub_2$, we call 'distal' the Ub unit connected through its C-terminus to K48 of the other Ub unit that is called 'proximal'. We define the 'proximal domain' and 'distal domain' binding sites on PLpro as S1 and S2, respectively, while the S1' site hangs off below the active site[7]. mISG15 is shown in cartoon representation and colored magenta. In analogy with the two Ubs in $Ub_2$, the N- and C-terminal UBL (ubiquitin-like) domains of ISG15 will be referred to as the distal and proximal UBLs, respectively.

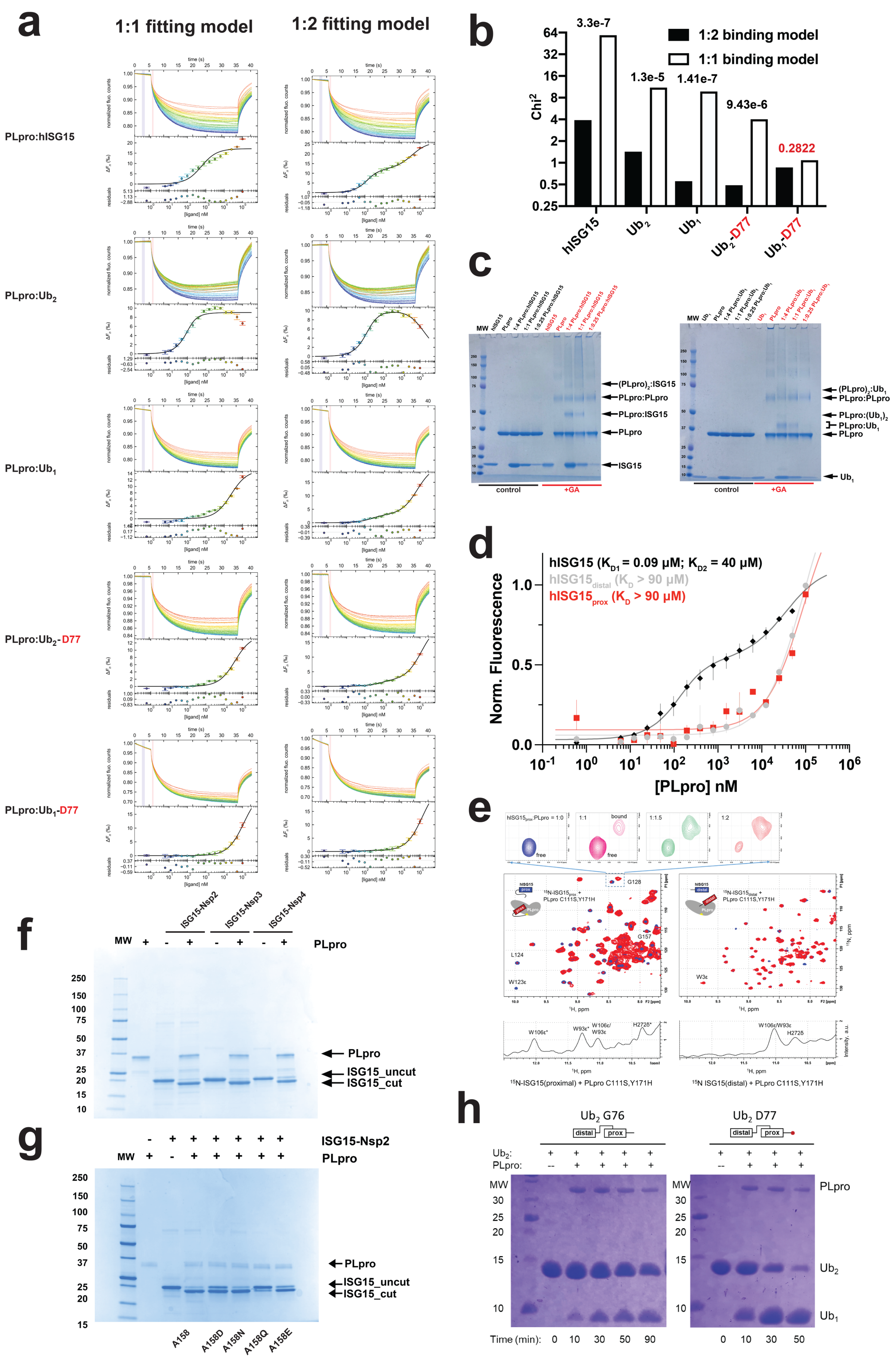


**Supplementary Figure 2. Fitting of 2:1 PLpro$^{CoV\text{-}2}$:substrate binding model data, individual hISG15 domain binding data, Nsp2-4 fusion constructs cleavage assays, and influence of C-terminal D77 on Ub$_2$ cleavage.** (**a**) Fitting the kinetics of binding between PLpro$^{CoV\text{-}2}$ to hISG15,

$Ub_2$, $Ub_1$, $Ub_2$ (D77) and $Ub_1$ (D77) to 1:1 (left) and 1:2 (right) stoichiometry binding models. Cumulative raw thermophoresis data for three independent titration experiments. The curves represent normalized fluorescence counts colored by concentration of PLpro from blue (low) to red (high) and plotted as a function of time. Fluorescence change ($\Delta F$) is plotted as a function of substrate concentration colored as above. Data is shown as averages with standard deviation across three independent experiments. Best fit is shown for 1:1 (left) and 1:2 (right) binding models. The variance between the average $\Delta F$ value and the fit is plotted below. The residuals are shown as average deviation from the fit across three experiments and colored as above. (**b**) Comparison of $\chi^2$ derived from the 1:1 and 1:2 fitting models for the $PLpro^{CoV-2}$ binding to hISG15, $Ub_2$, $Ub_1$, $Ub_2$ (D77) and $Ub_1$ (D77) binding data. The numbers above the bars are the probabilities that the observed improvement in the fit occurred by chance. Only for the $PLpro^{CoV-2}$:$Ub_1$ (D77) did the 1:2 model not statistically improve (Prob=0.2822). All other binding data were better explained by the 1:2 binding model with statistical significance. (**c**) Glutaraldehyde cross-linking (0.05%) of $PLpro^{CoV-2}$:hISG15 (left) or $PLpro^{CoV-2}$:$Ub_1$ (right) using 1:4, 1:1 and 1:0.25 ratios and resolved by SDS-PAGE. (**d**) MST binding curves for full-length hISG15 (black, 1:2 fitting model) and for the distal ($hISG15_{distal}$, gray, 1:1 fitting model) and proximal ($hISG15_{prox}$, red, 1:1 fitting model) UBLs binding to $PLpro^{CoV-2}$. Data is shown as an average of three replicates with range of measurements. (**e**) NMR-based binding experiment of $^{15}N$-labeled $hISG15_{prox}$ (left panel) and $hISG15_{distal}$ (right panel) UBL domains of hISG15 to unlabeled $PLpro^{CoV-2}$. Shown are 2-D $^{1}H$-$^{15}N$ SOFAST-HMQC spectra of each hISG15 UBL domain alone (blue) overlaid with the spectra (red) in the presence of 2-fold (left) or 4-fold (right) molar excess of unlabeled $PLpro^{CoV-2}$. Select residues are indicated; the residue numbers correspond to full-length hISG15: $hISG15_{distal}$ includes residues 2-82 and $hISG15_{prox}$ residues 81-157 of hISG15. Inset on top: zoom in on the spectral region containing free and bound signals of residue G128 in $^{15}N$-$hISG15_{prox}$, upon titration with 1x (magenta), 1.5x (green), and 2x (red) molar equivalents of $PLpro^{CoV-2}$. Shown on the bottom are fragments of $^{1}H$ 1-D spectra of the same samples as above (left: $^{15}N$-$ISG15_{prox}$+PLpro at 1:2; right: $^{15}N$-$ISG15_{distal}$+PLpro at 1:4), zoomed in on the region containing characteristic proton signals of $PLpro^{CoV-2}$'s indole NH group of tryptophans W93 and W106 and of imidazole NH of active-site histidine H272; the bound (marked with asterisk) and unbound signals are indicated. The observation of the bound (shifted) signals of these groups indicates that the C-terminus of $ISG15_{prox}$ binds the active site of PLpro (see also Figure 3 and Supplementary Figure 8). No such signals/shifts are detected for $ISG15_{distal}$. (**f**) $PLpro^{CoV-2}$ cleavage of hISG15-Nsp C-terminal fusions. SDS-PAGE gel reveals that PLpro cleaves off fragments of Nsp2 (LRGG↓AYTRYVDNNF), Nsp3 (LRGG↓APTKVTFGDD), and Nsp4 (LRGG↓KIVNNWLKQL) from hISG15-Nsp fusions linked to the recognition motif "LRGG". (**g**) $PLpro^{CoV-2}$ cleavage of hISG15-Nsp2 C-terminal fusions containing mutations at the first residue (Asn, Asp, Gln and Glu) (Nsp2, LRGG↓**A**YTRYVDNNF) after the "LRGG" motif and shown in bold. SDS-PAGE gel reveals that PLpro cleaves off fragments of Nsp2 (**X**YTRYVDNNF), (**h**) $PLpro^{CoV-2}$ cleavage of $Ub_2$ and $Ub_2$ (D77). SDS-PAGE gel reveals that PLpro cleaves $Ub_2$ (D77) more quickly than $Ub_2$ into $Ub_1$.

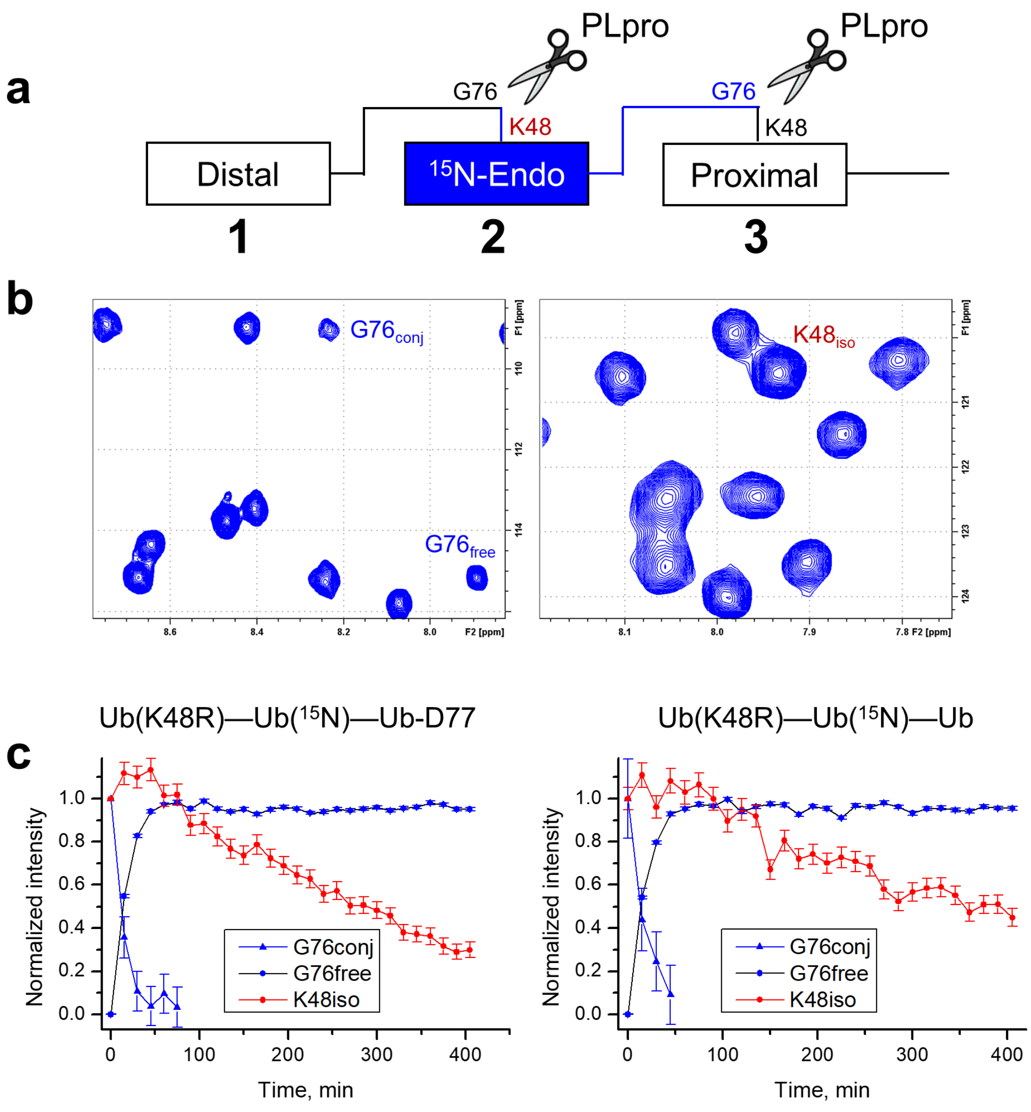


**Supplementary Figure 3. $PLpro^{CoV-2}$ cleavage of $Ub_3$ monitored by NMR.** (**a**) Cartoon representation of the $Ub_3$ constructs used (see also Figure 1g in main text). These $Ub_3$ constructs provided a unique opportunity to monitor in real time by NMR and directly compare cleavage of both isopeptide bonds simultaneously. (**b**) Fragments of $^1H$-$^{15}N$ SOFAST-HMQC NMR spectrum of $Ub_3$ shown in (a) zooming on the signals of G76 and the isopeptide-linked K48 of the $^{15}N$-labeled endo Ub. (**c**) Normalized intensities of NMR signals of G76 (free and conjugated) and of the side chain NH of isopeptide-bonded K48 ($K48_{iso}$) after mixing the $Ub_3$ with $PLpro^{CoV-2}$. (Left) the same $Ub_3$ construct as used for the MS studies shown in Figure 1 (main text). (Right) $Ub_3$ with WT Ub used as the proximal Ub. $PLpro^{CoV-2}$ was added in 1:10000 molar ratio. The intensities were normalized to either the first data point (for $G76_{conj}$ and $K48_{iso}$) or the last (for $G76_{free}$). The error bars represent uncertainties in the normalized peak intensities calculated based on the measured noise in the NMR spectra. Note that the disappearance of $G76_{conj}$ and the concomitant emergence of $G76_{free}$ signals happen significantly faster than the disappearance of the $K48_{iso}$ signal, indicating that the proximal Ub (unit 3) gets cleaved first, while the isopeptide bond between the distal and the endo Ubs (units 1 and 2) is cleaved much slower, in agreement with the MS data. A slight increase in the NMR signal intensity of $K48_{iso}$ at the early stage of the cleavage reaction reflects faster tumbling (due to smaller size) of the resulting Ub chain (containing the $^{15}$N-labeled Ub) when the proximal Ub is cleaved off.

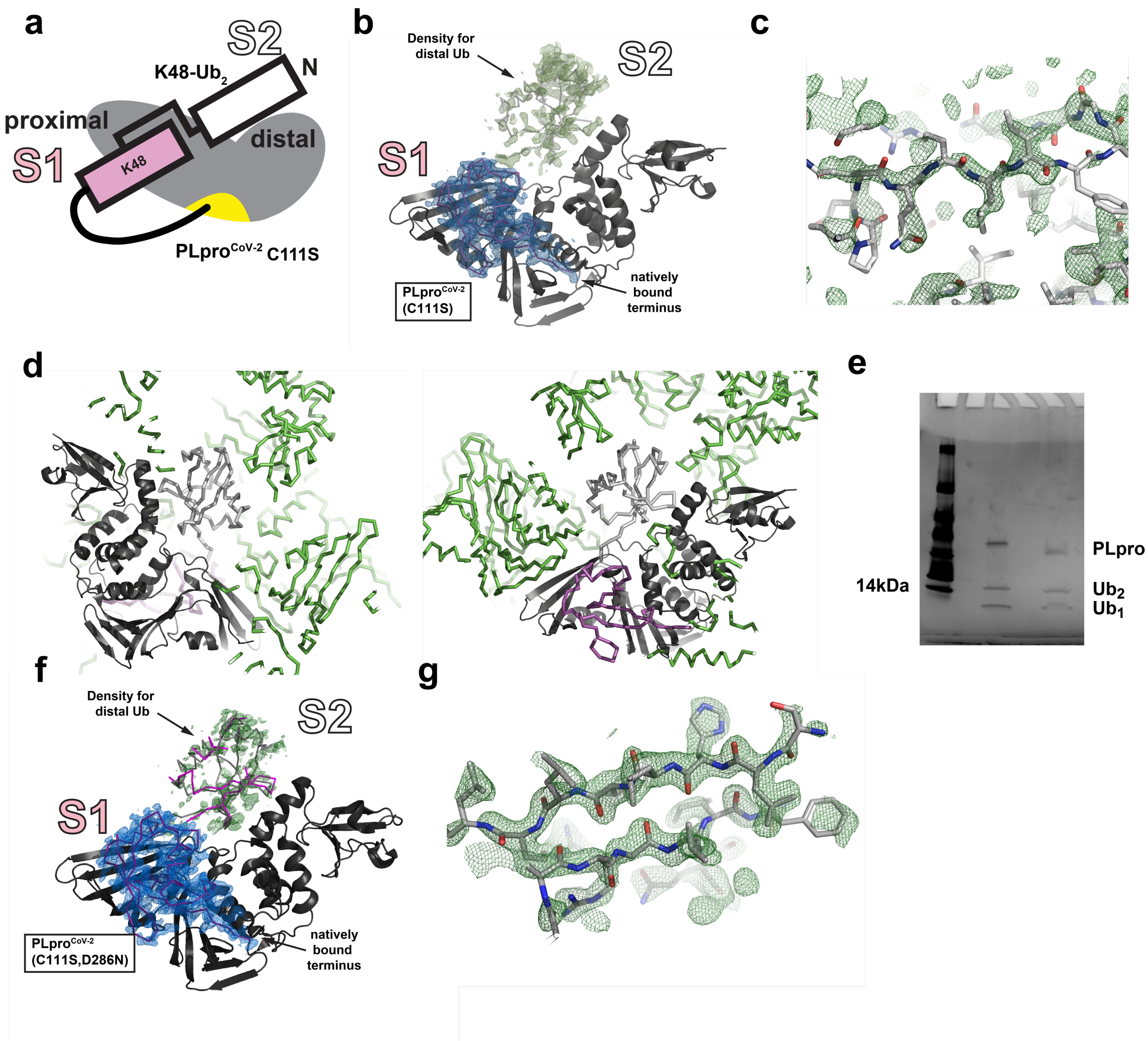


**Supplementary Figure 4. Strong electron density in the proximal domain for PLpro$^{CoV-2}$: K48-Ub$_2$ complex.** (**a**) Schematic illustration for PLpro$^{CoV-2}$:K48-Ub$_2$ complex. (**b**) Structure of PLpro$^{CoV-2}$ in complex with K48-Ub$_2$ overlayed with obtained electron density maps for K48-Ub$_2$ substrate. PLpro is shown in cartoon representation and colored dark gray. K48-Ub$_2$ is shown as ribbons and colored magenta and light gray for the proximal and distal Ubs, respectively. The distal Ub model was obtained from the structure of SARS-CoV-1 PLpro bound to a K48-linked Ub$_2$ (PDB id: 5E6J) superposed on our SARS-CoV-2 PLpro in complex with a K48-linked Ub$_2$. Arrows show strong electron density map fragment for C-terminus of proximal domain of K48-Ub$_2$ and observed indication of helical density in its distal domain. The $2F_o - F_c$ electron density weighted map for the proximal Ub is shown as a blue mesh, contoured at $1.4\sigma$. The $F_o - F_c$ electron density weighted difference map for the distal Ub area is shown as a green mesh, contoured at 2σ. (**c**) Close-up of area corresponding to the distal Ub not modelled in the structure showing presence of electron density resembling amino acid chain with side chains corresponding to Ub sequence. The sequence in center corresponding to distal Ub residues 41-44 (PDB id: 5E6J) is shown in gray with oxygens and nitrogen atoms in red and blue, respectively. The $F_o - F_c$ electron density weighted difference map is shown as a green mesh, contoured at 1.6σ. The 40-46 Ub fragment was moved to the density by Rigid Body Fit Zone in Coot program without any side-chain adjustments. The shift of Cα atoms was in 1.3-2.3 Å range. (**d**) The distal Ub not modelled in the structure occupies the "empty" space in $PLpro^{CoV-2}$:$Ub_2$ crystal model. PLpro is shown in cartoon representation and colored dark gray. K48-$Ub_2$ is shown as ribbons and colored magenta and light gray for the proximal and distal Ubs, respectively. The neighbor molecules of $PLpro^{CoV-2}$:$Ub_{prox}$ are shown as green ribbons. (**e**) SDS-PAGE gel of $PLpro^{CoV-2}$ (C111S):$Ub_2$ crystals used for data collection. PLpro, $Ub_2$ and $Ub_1$ are highlighted. (**f**) Structure of $PLpro^{CoV-2}$ (C111S,D286N) in complex with K48-$Ub_2$ overlayed with obtained electron density maps for K48-$Ub_2$ substrate. PLpro is shown in cartoon representation and colored dark gray. K48-$Ub_2$ is shown as ribbons and colored magenta, and overlaid distal Ub (from PDB id: 5E6J) in gray. Arrows show strong electron density map fragment for C-terminus of proximal domain of K48-$Ub_2$ and observed indication of helical density in its distal domain. The $2F_o - F_c$ electron density weighted map for the proximal Ub is shown as a blue mesh, contoured at 1.5*σ*. The $F_o - F_c$ electron density weighted difference map for the distal Ub area is shown as a green mesh, contoured at 2σ. (**g**) Close-up of area corresponding to the distal Ub not modelled in the structure showing presence of electron density resembling amino acid chain with side chains corresponding to Ub sequence. The sequence in center corresponding to distal Ub residues 40-45 (lower β-strand) and 66-73 (upper β-strand) (PDB id: 5E6J) is shown in gray with oxygens and nitrogen atoms in red and blue, respectively. The $2F_o - F_c$ electron density weighted difference map is shown as a green mesh, contoured at 0.9σ.

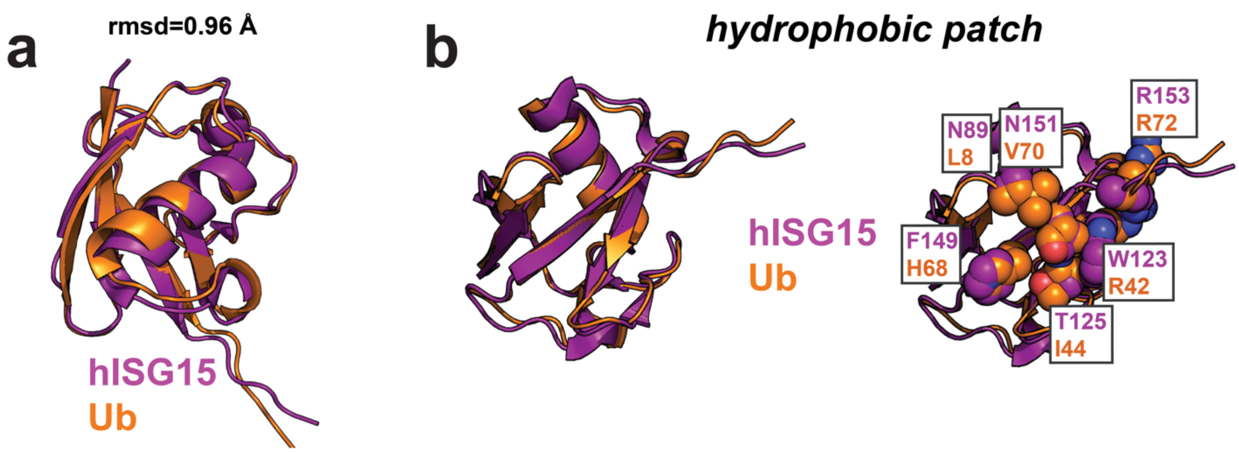


**Supplementary Figure 5. Comparison of hISG15 and Ub$_1$ bound to PLpro$^{CoV-2}$.** (**a**) Structural alignment of proximal domain of hISG15 to Ub shows high similarity with a Cα rmsd of 0.96 Å. Both, hISG15 and Ub, are shown in cartoon representation. hISG15 is colored magenta and Ub is orange. (**b**) Comparison of binding surface for proximal hISG15 and Ub plus additional contacts involved in binding to PLpro. Key residues are shown in spacefill and are colored as in (a).

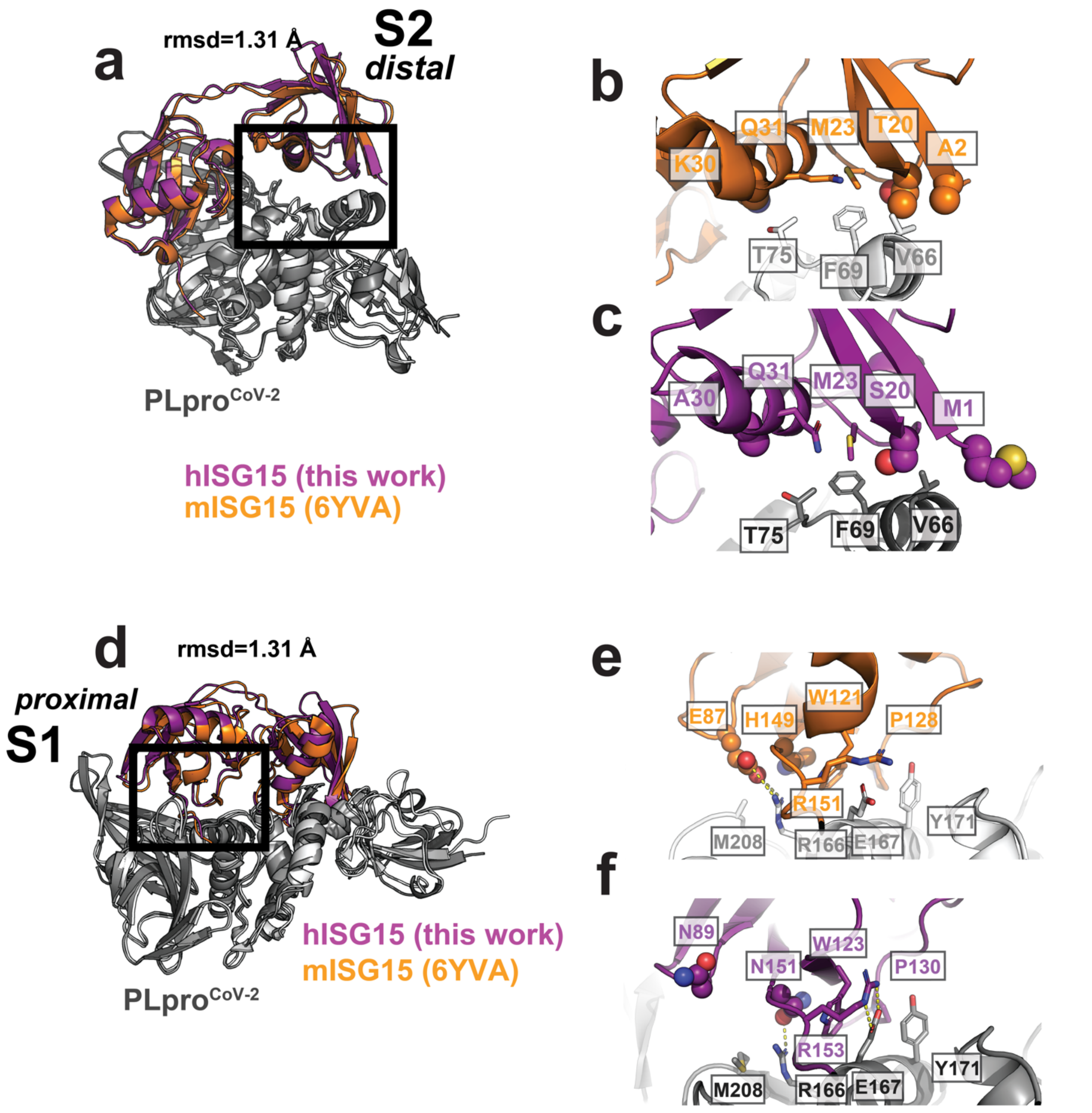


**Supplementary Figure 6. Comparison of contacts between hISG15 and mISG15 bound to PLpro$^{CoV-2}$.** (**a**) Structural overlay of PLpro$^{CoV-2}$:hISG15 (gray and magenta) from this study (PDB id: 7RBS) and PLpro$^{CoV-2}$:mISG15 (white and orange) from previously published studies (PDB id: 6VYA) focusing on the distal site (S2). (**b, c**) Zoom in of boxed contacts from (a) from PLpro to the distal (S2) domain of (**b**) mISG15 or (**c**) hISG15. Interface residues are shown in sticks and colored as in (a). Amino acids that vary between mISG15 and hISG15 are shown in spheres. Interface amino acids are labeled. (**d**) Structural overlay of PLpro$^{CoV-2}$:hISG15 and PLpro$^{CoV-2}$:mISG15 focusing on the proximal site (S1) and colored as in (a). (**e, f**) Zoom in of the boxed contacts from (d) from PLpro to the proximal domain of (**e**) mISG15 or (**f**) hISG15. Interface residues are shown in sticks and colored as in (d). Amino acids that vary between mISG15 and hISG15 are shown in spheres. Interface amino acids are labeled. Hydrogen bonds are shown as dashed yellow lines.

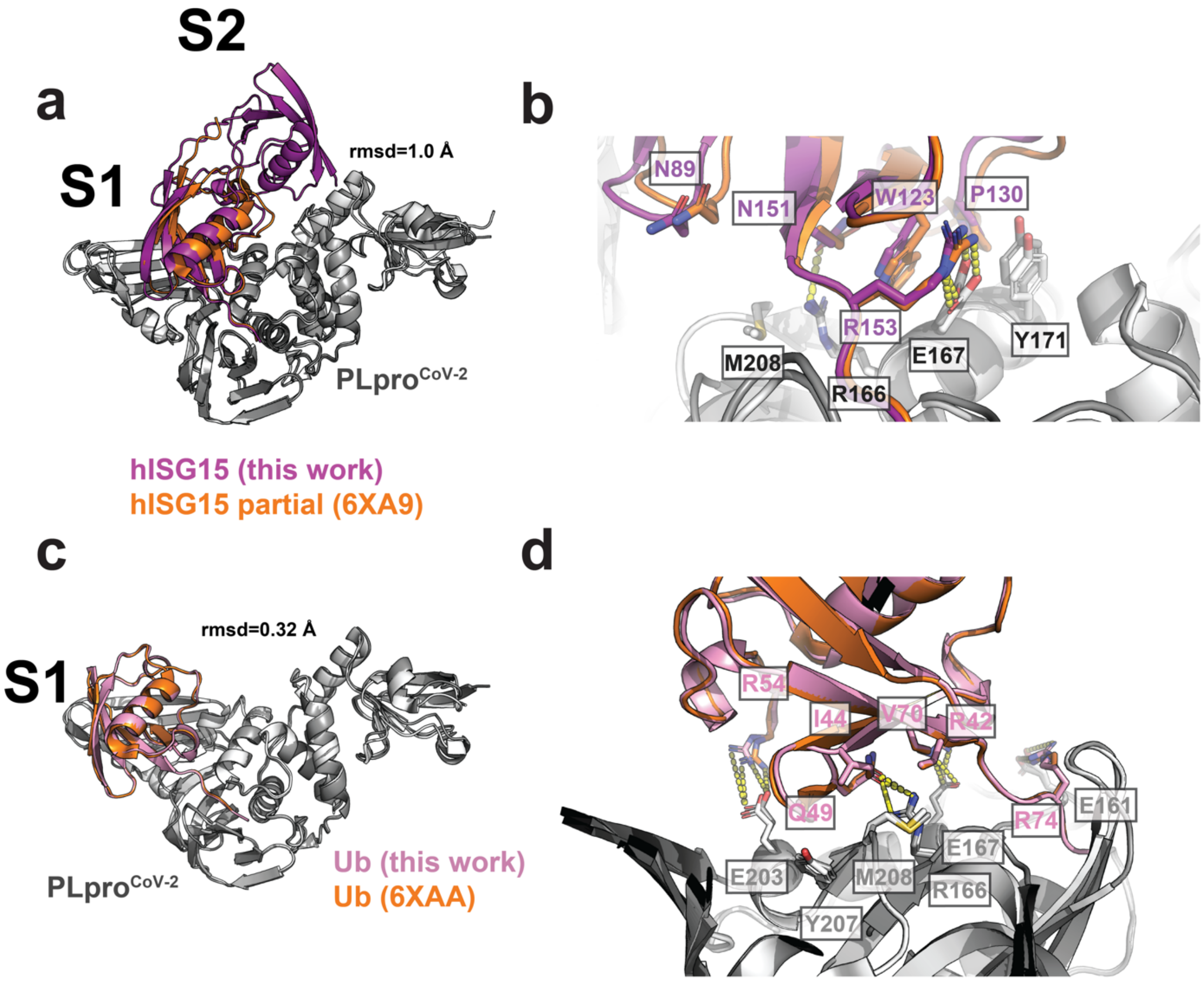


**Supplementary Figure 7. Comparison of partial hISG15 and full hISG15 bound to PLpro$^{CoV-2}$.** (**a**) Structural overlay of PLpro$^{CoV-2}$ in complex with hISG15 from this work (PDB id: 7RBS) and from previously published studies with a partial hISG15 (PDB id: 6XA9) shows a Cα rmsd of 1.0 Å. PLpro and hISG15 are shown in cartoon representation. PLpro and hISG15 from this paper are colored gray and magenta, respectively. PLpro and partial hISG15 from previous studies are colored white and orange, respectively. (**b**) Zoom in of the interface comparing the two structures and colored as in (a). Interface residues are labeled and are shown as sticks. Hydrogen bonds are shown as dashed yellow lines. (**c**) Overlay of $PLpro^{CoV-2}$ in complex with proximal domain of K48-$Ub_2$ from this paper (PDB id: 7RBR) and from previously published studies (PDB id: 6XAA) shows a C$\alpha$ rmsd of 0.32 Å. PLpro and Ub are shown in cartoon representation. PLpro and proximal domain of Ub from this study is colored gray and pink. PLpro and proximal domain of Ub from previous study is colored in white and orange. (**d**) Zoom in of the interface comparing the two structures and colored as in (c). Interface residues are labeled and are shown as sticks. Hydrogen bonds are shown as dashed yellow lines.

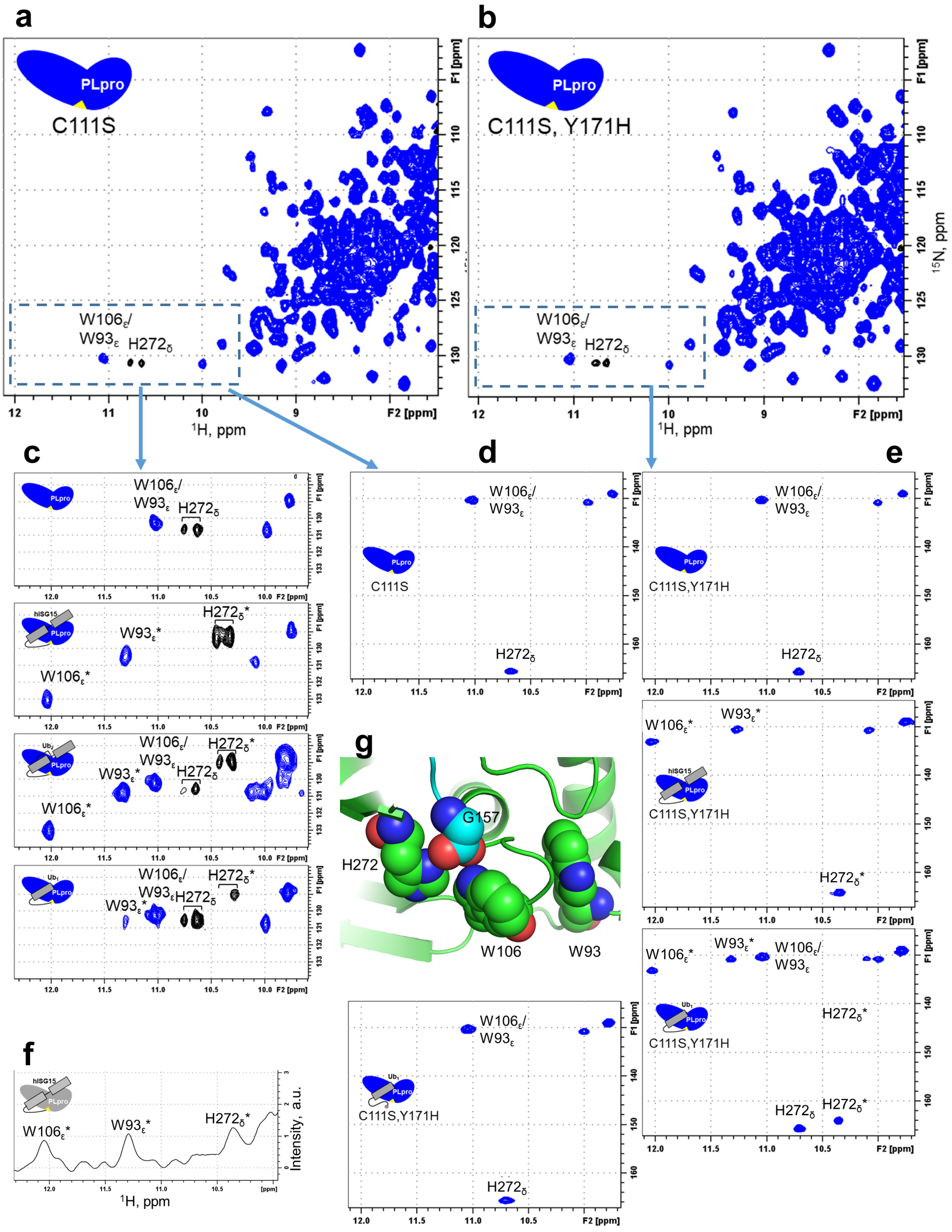

a
b
C111S
C111S, Y171H
PLpro
W106ε/
W93ε
H272δ
¹H, ppm
¹⁵N, ppm
F1 [ppm]
F2 [ppm]
c
d
e
f
g
hISG15
Ub₂
Ub₁
W93ε*
W106ε*
H272δ*
C111S,Y171H
G157
H272
W106
W93
Intensity, a.u.

**Supplementary Figure 8. NMR characterization and comparison of substrate binding to PLpro$^{CoV-2}$ C111S and C111S,Y171H variants.** (**a, b**) Comparison of $^{1}H$-$^{15}N$ SOFAST-HMQC spectra of $^{15}N$-labeled PLpro$^{CoV-2}$ (C111S) (**a**) and PLpro$^{CoV-2}$ (C111S,Y171H) (**b**). The region containing side chain NH signals of W93, W106, and H272 is marked by a dashed rectangle. Positive signals are blue, negative are black. The histidine imidazole NH group has $^{15}N$ chemical shift of 165.5 ppm. Therefore, its signal is aliased (folded in) in this spectrum, resulting in negative intensity. It appears as a doublet (consistent with $^{1}J \sim 94$ Hz) because of the inefficient decoupling of $^{15}N$ during $^{1}H$ acquisition time due to off-resonance effects. See also panels (d and e) where these effects were removed. The crosspeaks corresponding to tryptophan indole and histidine imidazole NH groups are indicated as $W_{\varepsilon}$ and $H_{\delta}$, respectively. (**c**) Zoom in on the spectral region containing Trp/His signals (rectangle in panel (a)) for $^{15}N$-PLpro (C111S) alone and upon addition of 1.5 molar equivalents of unlabeled ISG15 or K48-Ub$_2$ or of 7 molar equivalents of WT Ub$_1$. The unbound ($W_{\varepsilon}$ and $H_{\delta}$) and shifted ($W_{\varepsilon}$* and $H_{\delta}$*) side chain tryptophan and histidine signals are indicated. The coloring of spectra is the same as in (a, b). (**d**) Zoom in on the region of $^{1}H$-$^{15}N$ SOFAST-HMQC spectrum of $^{15}N$-PLpro (C111S) acquired with the $^{15}N$ spectral width of 70 ppm that covers the NH signals from both amides and the side chains of Trp and His, eliminating aliasing and splitting of the His signal. The coloring of signals is the same as in (a, b). (**e**) Zoom on the region of $^{1}H$-$^{15}N$ SOFAST-HMQC spectra of $^{15}N$-PLpro (C111S,Y171H) alone and in the presence of the indicated substrates, acquired with the $^{15}N$ spectral width of 90 ppm that covers the NH signals from both amides and from the side chains of Trp and His, thus eliminating aliasing and splitting of His signals. The unbound ($W_{\varepsilon}$ and $H_{\delta}$) and shifted ($W_{\varepsilon}$* and $H_{\delta}$*) side chain tryptophan and histidine NH signals are indicated. The coloring of signals is the same as in (a, b). (**f**) Fragment of $^{1}H$ 1-D spectrum of a $^{15}N$-hISG15+(unlabeled) PLpro$^{CoV-2}$ (C111S) sample (at 1:1 ratio), zoomed in on the region containing characteristic proton signals of PLpro$^{CoV-2}$'s indole NH group of tryptophans W93 and W106 and of imidazole NH of active-site histidine H272, as indicated, corresponding to the hISG15-bound state of PLpro. (**g**) Fragment of the PLpro$^{CoV-2}$:hISG15 structure (PDB id: 7RBS) zooming on the active site of the enzyme to illustrate a close location of the C-terminus of hISG15 (G157) to PLpro$^{CoV-2}$ residues W93, W106, and H272 (shown as spheres). PLpro is colored green, hISG15 is cyan. Carboxylic oxygens of the C-terminal G157 form polar contacts with $N_{\delta}$ of H272 and $N_{\varepsilon}$ of W106. The contacts between G76 of the proximal Ub of K48-Ub$_2$ and PLpro$^{CoV-2}$ residues W93, W106, and H272 are very similar (PDB id: 7RBS).

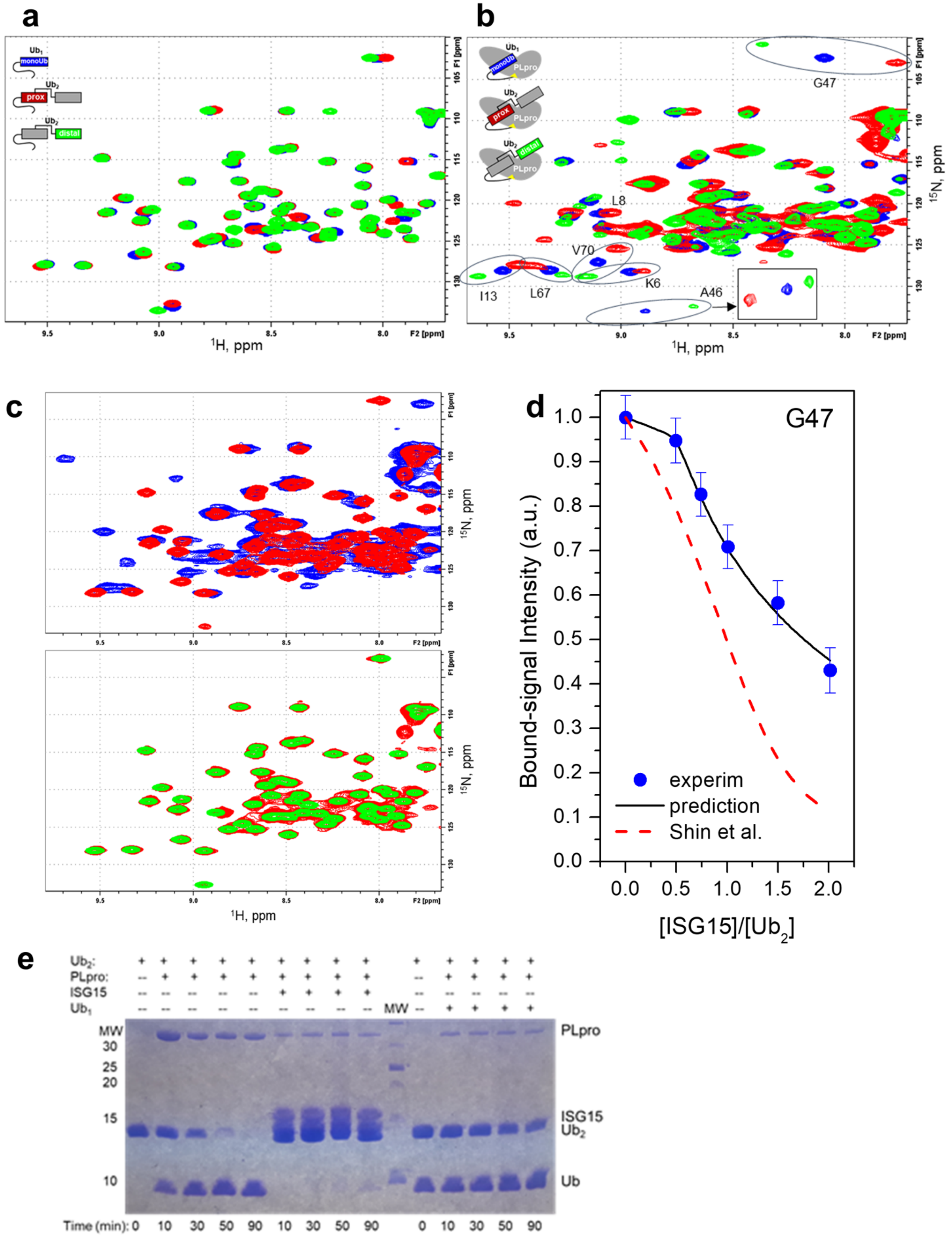
a
b
c
d
e
Ub1
monoUb
Ub2
prox
distal
PLpro
G47
L8
V70
K6
I13
L67
A46
1H, ppm
15N, ppm
F2 [ppm]
F1 [ppm]
G47
Bound-signal Intensity (a.u.)
[ISG15]/[Ub2]
experim
prediction
Shin et al.
Ub2:
PLpro:
ISG15
Ub1
MW
PLpro
ISG15
Ub2
Ub
Time (min): 0 10 30 50 90 10 30 50 90 0 10 30 50 90

**Supplementary Figure 9. Characterization of substrate binding to PLpro$^{CoV-2}$-C111S and of substrate competition for binding and cleavage.** (**a, b**) overlay of $^{1}H$-$^{15}N$ SOFAST-HMQC spectra of $Ub_1$ (blue) and of the proximal (red) and distal (green) Ubs in K48-$Ub_2$, free in solution (**a**) and in complex with PLpro$^{CoV-2}$-C111S (**b**) (see also Fig. 3). Examples illustrating the striking difference in the directions of signal shifts in the distal and proximal Ubs of several residues are circled. (**c**) NMR analysis of the competition between K48-$Ub_2$ and hISG15 for PLpro binding. Shown is an overlay of $^{1}H$-$^{15}N$ SOFAST-HMQC spectra of the $^{15}N$-labeled proximal Ub in K48-$Ub_2$ alone in solution (green), in complex with 1.5 molar equivalents of unlabeled PLpro$^{CoV-2}$-C111S (blue) and upon subsequent addition of 2 molar equivalents of unlabeled hISG15 (red). The spectra show pair-wise comparison. The red contour levels are selected to emphasize the strong signals, for better comparison with the spectra of $Ub_2$ alone. (**d**) Intensity of the PLpro-bound signal of G47 (from the binding competition assay as a function of [hISG15]:[$Ub_2$] molar ratio (dots, normalized to the initial intensity) and the predicted molar fraction of bound $Ub_2$ (black line) using the $K_{d1}$ values obtained in this work (see Fig. 3j). The dashed red line shows the predicted molar fraction of bound $Ub_2$ using the $K_d$ values reported by Shin et al. for hISG15 and triazole-linked $Ub_2$[8]. These predictions used equations describing competitive binding of two different ligands to the same binding site[9]. (**e**) SDS-PAGE gels showing the inhibitory effect of hISG15 (middle lanes) and $Ub_1$ (right lanes) on disassembly of K48-linked $Ub_2$ by PLpro$^{CoV-2}$ (see also Fig. 3k).

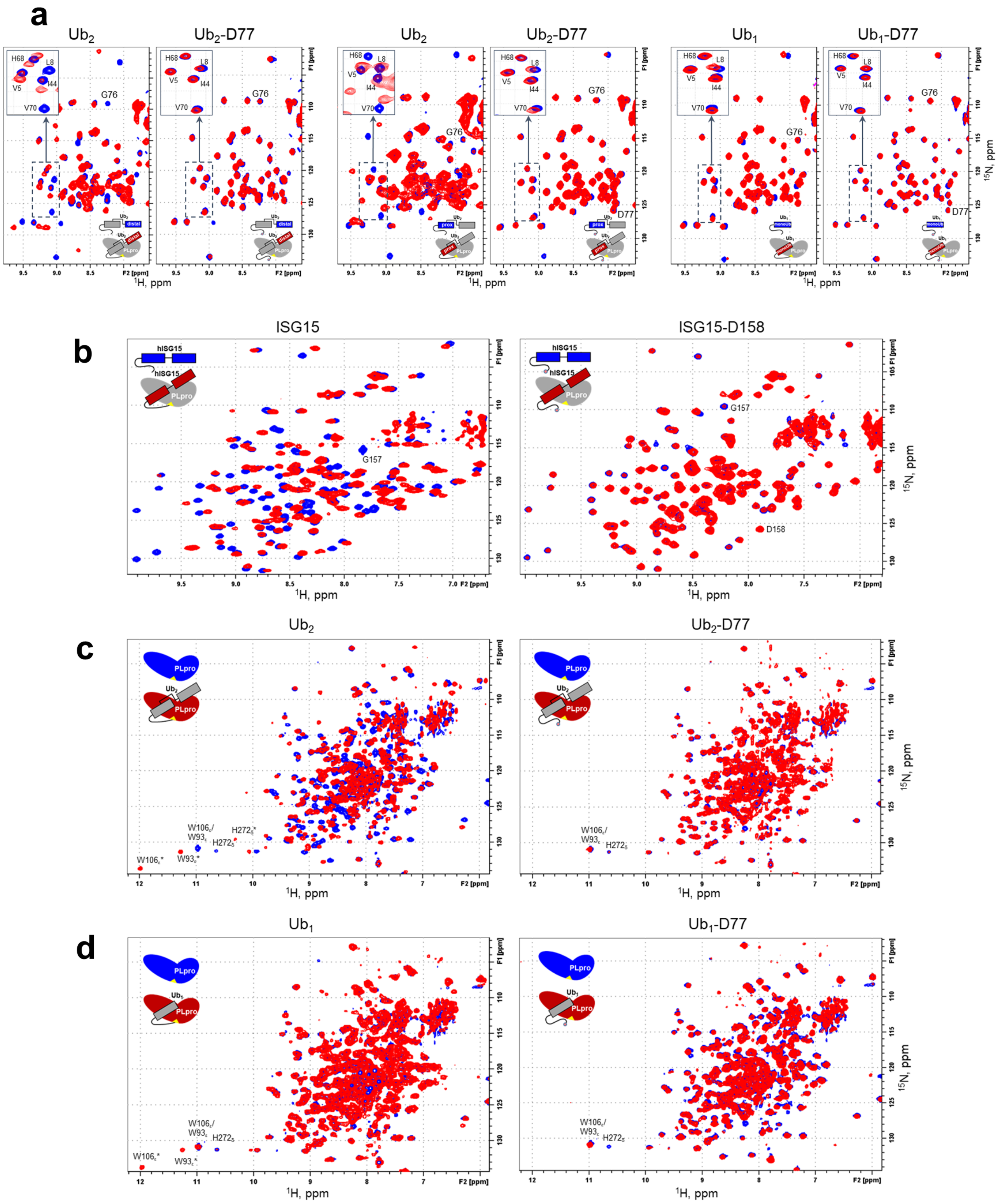


**Supplementary Figure 10. NMR characterization of the effect of addition of a C-terminal aspartate on interactions of hISG15, K48-Ub$_2$, and Ub$_1$ with PLpro$^{CoV\text{-}2}$.** (**a**) Overlay of $^1$H-$^{15}$N spectra of $^{15}$N-labeled distal and proximal Ubs in $Ub_2$ and $Ub_1$ alone (blue) and upon addition of PLpro$^{CoV-2}$-C111S. In each pair, the spectrum on the right corresponds to the presence of D77 at the C-terminus of $Ub_2$ or $Ub_1$. PLpro was added in 1:1 (for $Ub_2$) or 2:1 (for $Ub_1$) molar ratio for wild-type proteins and in 2:1 for the D77-extended constructs. Signals of select residues are indicated. Insets zoom on the spectral region containing signals of the hydrophobic patch residues L8, I44, H68, V70, as well as V5. (**b**) Overlay of $^1$H-$^{15}$N spectra of $^{15}$N-labeled hISG15 alone (blue) and upon addition of PLpro$^{CoV-2}$-C111S (left) or PLpro$^{CoV-2}$-C111S,Y171H (right). The spectra on the left (TROSY) correspond to WT hISG15, those on the right (SOFAST-HMQC) are for hISG15-D158. The signals corresponding to C-terminal G157 (hISG15) and G157 and D158 (hISG15-D158) are indicated. PLpro was added in ~1 molar equivalent to hISG15 and 1.5 molar equivalents to hISG15-D158. (**c**) Overlay of $^1$H-$^{15}$N TROSY spectra of $^{15}$N-labeled PLpro$^{CoV-2}$-C111S,Y171H alone (blue) and upon addition of K48-$Ub_2$ (left) or K48-$Ub_2$-D77 (right). The signals corresponding to side chain NHs of W93, W106, and H272 are marked as $W_\varepsilon$ and $H_\delta$ for the unbound signals and $W_\varepsilon$* and $H_\delta$* for the (shifted) bound signals. (**d**) Overlay of $^1$H-$^{15}$N TROSY spectra of $^{15}$N-labeled PLpro$^{CoV-2}$-C111S,Y171H alone (blue) and upon addition of 8 molar equivalents of WT $Ub_1$ (left) or $Ub_1$-D77 (right). The signals corresponding to side chain NHs of W93, W106, and H272 are marked as $W_\varepsilon$ and $H_\delta$ for the unbound signals and $W_\varepsilon$* and $H_\delta$* for the (shifted) bound signals.

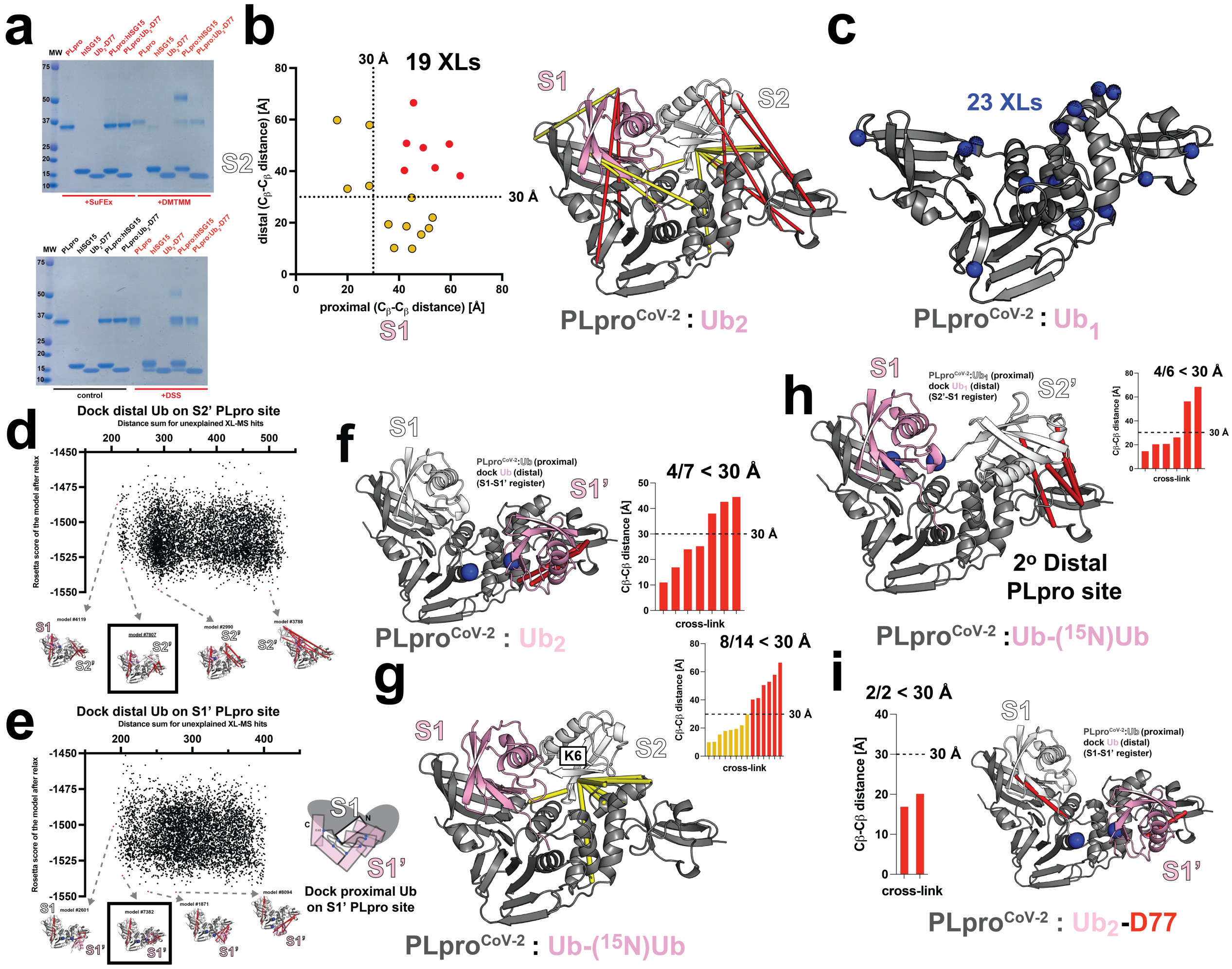


**Supplementary Figure 11. Intermolecular cross-links from XL-MS analysis of PLpro$^{CoV-2}$ complexes.** (**a**) Cross-linked PLpro$^{CoV-2}$:hISG15 and PLpro$^{CoV-2}$:Ub$_2$-D77 complexes reveal formation of covalent heterodimer complex bands (red) compared to control reactions (untreated and PLpro$^{CoV-2}$ or substrate alone) by SDS-PAGE. (**b**) Intermolecular cross-links sites (19 pairs) mapped onto proximal or distal Ub binding modes on PLpro$^{CoV-2}$. Cross-link pair distances between Cβ-Cβ below 30 Å or above 30 Å are colored in yellow and red, respectively. Calculated proximal and distal distances for cross-link pairs are plotted on a scatter plot. Cross-links are mapped onto structure of PLpro:Ub$_2$ and colored yellow or red based on distance. Structures are shown in cartoon and colored as previously. Cross-links are shown as lines. (**c**) Cross-links derived from PLpro$^{CoV-2}$:Ub$_1$ complexes (23 pairs). Mapping Ub contact points on PLpro$^{CoV-2}$ derived from heterotypic cross-links pairs. PLpro$^{CoV-2}$ is shown as cartoon (gray), cross-link sites are shown as spheres (blue). (**d, e**) Docking Ub onto PLpro$^{CoV-2}$:Ub$_{prox}$ (S1) to identify alternative S2 site, termed S2' (**d**) and S1' (**e**) sites employing a geometric restraint that links K48 from proximal Ub to C-terminus of the distal Ub but with two different binding registers: (1) proximal Ub in S1 binding site and distal sampling alternative S2' sites (d) or (2) distal Ub in S1 with the proximal Ub sampling alternative S1' sites (e). Energy of complex for over 5000 models was compared to the sum of distances for 7 unexplained cross-links from (b, in red). Below representative structures that sample diverse geometries of the docked Ub that are compatible or incompatible with cross-link distances. Structures are shown in cartoon and colored as before. Ub domains are shown as cartoon on S2'/S1 (d) or S1/S1' binding sites (e). Cross-links are shown as red lines. (e, right) Schematic for illustrating docking scheme, Ub docked onto PLpro$^{CoV-2}$:Ub$_{prox}$ into the S1' site. (**f**) Seven previously unexplained heterotypic cross-links identified from a PLpro$^{CoV-2}$:Ub$_2$ complex. Cross-links between Ub and PLpro$^{CoV-2}$'s thumb and UBL domains are mapped onto a docked model of PLpro$^{CoV-2}$:Ub with Ub placed in the S1' site based on the K48-Cterm linkage. Structures are shown as cartoons and colored as previously. The cross-links pairs shown as red lines. Bar plot of calculated distances for 7 cross-links and colored in red. A dashed line denotes the 30 Å threshold. (**g**) Fourteen heterotypic cross-links identified from a PLpro$^{CoV-2}$:Ub-($^{15}$N)Ub complex between the protease and the light Ub peptides. Cross-links are mapped onto a model of PLpro$^{CoV-2}$ with the canonical (i.e., as found in the crystal structure) Ub$_2$ binding mode in S1/S2 binding sites. Structures are shown as cartoons and colored as previously. The cross-links pairs shown as yellow lines. Bar plot of calculated distances for 14 cross-links and colored in yellow (< 30 Å threshold) or red (> 30 Å threshold). A dashed line denotes the 30 Å threshold. (**h**) Six previously unexplained heterotypic cross-links identified from (g, red). Cross-links are mapped onto a docked model of PLpro$^{CoV-2}$:Ub$_{prox}$ with Ub placed in the S2' based K48-Cterm linkage geometry. Structures are shown as cartoons and colored as previously. The cross-links pairs shown as red lines. Bar plot of calculated distances for 6 cross-links and colored in red. A dashed line denotes the 30 Å threshold. (**i**) Two heterotypic cross-links identified from the PLpro$^{CoV-2}$:Ub$_2$-D77 complex. Cross-links are mapped onto a docked model of PLpro$^{CoV-2}$:Ub$_{prox}$ with Ub placed in the S1' based K48-Cterm linkage geometry. Structures are shown as cartoons and colored as previously. The cross-links pairs shown as red lines. Bar plot of calculated distances for 2 cross-links and colored in red. A dashed line denotes the 30 Å threshold.

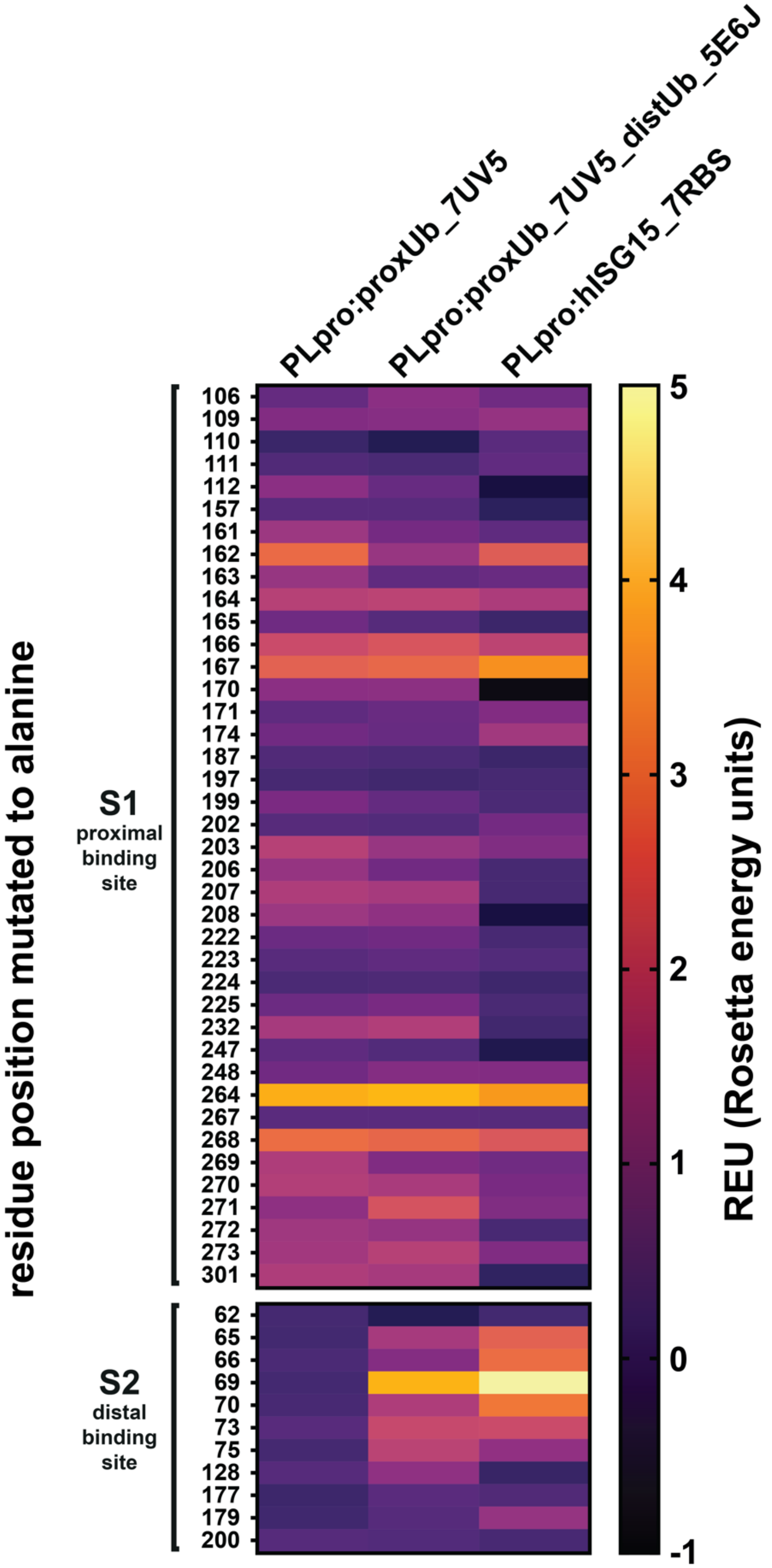
PLpro:proxUb_7UV5
PLpro:proxUb_7UV5_distUb_5E6J
PLpro:hISG15_7RBS
residue position mutated to alanine
S1
proximal
binding
site
S2
distal
binding
site
106
109
110
111
112
157
161
162
163
164
165
166
167
170
171
174
187
197
199
202
203
206
207
208
222
223
224
225
232
247
248
264
267
268
269
270
271
272
273
301
62
65
66
69
70
73
75
128
177
179
200
5
4
3
2
1
0
-1
REU (Rosetta energy units)

**Supplementary Figure 12. Interactions stabilizing hotspot residues in $PLpro^{CoV-2}$ in complex with substrates.** Heat-map results of $\Delta\Delta G_{binding}$ calculations of *in silico* alanine scan for $PLpro^{CoV-2}$ in complex with $Ub_1$ (based on PDB id: 7UV5), $Ub_2$ (distal Ub modeled based on PDB id: 5E6J orientation) and hISG15 (PDB id: 7RBS). Row titles represent positions of residues on the S1 (proximal) and S2 (distal) interaction surface residues are labeled. The heat-map is colored from black to yellow.

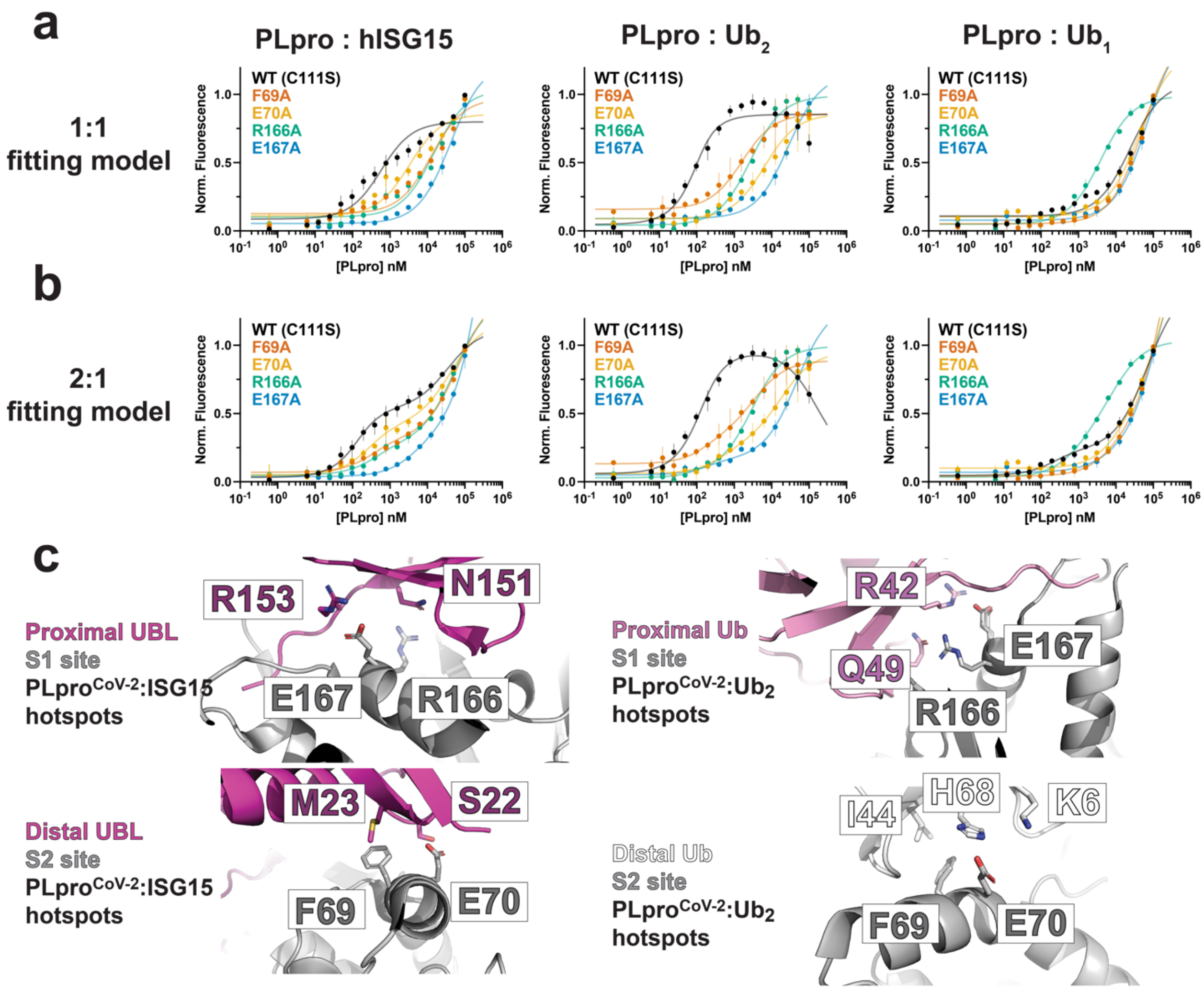


**Supplementary Figure 13. MST measurement of PLpro$^{CoV-2}$ mutants binding to hISG15, $Ub_2$ and $Ub_1$.** MST titrations are fitted to a 1:1 (**a**) and 1:2 (**b**) binding model using PALMIST software. Data is shown as an average of three replicates with range of measurements. (**c**) Illustration of the hotspot interactions surrounding E167, R166, F69 and E70 in PLpro$^{CoV-2}$:hISG15 (left) and PLpro$^{CoV-2}$:$Ub_2$ complexes (right). Structures are shown in cartoon representation and the residues interacting with the hotspots are shows as sticks. PLpro is colored in gray. Proximal and distal UBLs of hISG15 are colored in magenta. Proximal and distal Ubs are colored pink and white, respectively.

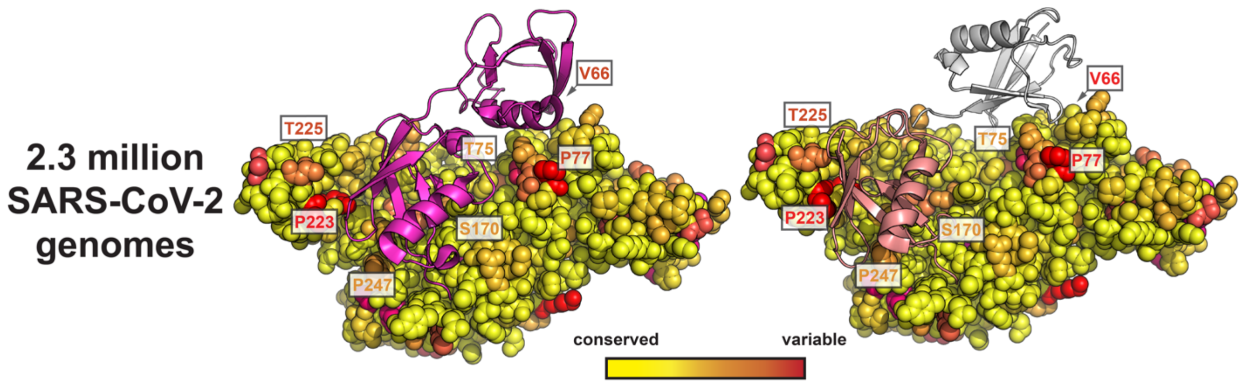


**Supplementary Figure 14. Evolutionary analysis of PLpro across SARS-CoV-2 variants.** Sequence variation across 2.3 million PLpro$^{CoV-2}$ isolates reveals variation in both the proximal and distal binding sites. PLpro is shown in space fill representation and is color-coded according to conservation from yellow to red. Sites of variation are labeled. K48-linked $Ub_2$ and human ISG15 binding modes are shown for comparison and colored as in Supplementary Fig. 1.
